# Loss of interruption in the *HTT* CAG repeat is associated with increased somatic expansion and loss of medium spiny neurons in HD

**DOI:** 10.64898/2026.03.02.709181

**Authors:** Chris Kay, Jessica Dawson, Kert Mätlik, Hailey Findlay Black, Emily Harvey, Stephanie Bortnick, Kyla Javier, Christina Buchanan, Tanushri Soomarooah, Inês Caldeira Brás, Glen Sequiera, Mahmoud Pouladi, Larissa Arning, Hoa HP Nguyen, Richard Roxburgh, Maurice Curtis, Richard LM Faull, Nathaniel Heintz, Michael R. Hayden

**Author notes:** These authors contributed equally to this manuscript (Chris Kay, Jessica Dawson).

## Abstract

Synonymous loss of interruption variants in the expanded CAG repeat sequence of Huntingtin (*HTT*) accelerate the clinical onset and progression of Huntington disease (HD). Medium spiny neurons (MSNs) are gradually lost in HD and undergo selective somatic CAG expansion, but it is unclear how CAG expansion in MSNs relates to HD pathology. Here, we show that MSNs with large (111-150 CAG) and very large (>150 CAG) somatic expansions are rare in early manifest HD, but accumulate in proportion with duration of disease and inherited CAG repeat length. In patients with the deleterious CAG-CCG loss-of-interruption (CAG-CCG LOI) modifier, the proportion of MSNs with large and very large expansions is increased ∼5-fold despite reduced small somatic expansions in blood, and direct caudate MSN counts are reduced. Our findings suggest that increased somatic CAG expansion contributes to accelerated striatal MSN pathology and hastened onset of HD, but that MSNs with very large genomic CAG expansions can persist among surviving neurons of the HD brain.

## Introduction

Huntington disease (HD) is a fatal repeat expansion disorder characterized by progressive motor dysfunction, psychiatric disturbances, cognitive decline, and selective degeneration of the striatum^1,2^. HD is caused by 36 or more uninterrupted CAG repeats in exon 1 of the Huntingtin gene (*HTT*), and the number of inherited CAG repeats is the primary determinant of the age at which symptoms first emerge^3–7^. Although *HTT* is ubiquitously expressed, the CAG repeat expands variably across tissues and cell types, with the greatest somatic expansion occurring in striatal neurons that are selectively lost in HD^8–13^. Somatic expansion of the CAG repeat increases in blood and tissues of HD patients with inherited CAG repeat length and age, suggesting a role for somatic expansion in the CAG- and age-dependent onset of HD^14–17^.

Approximately a dozen genetic modifiers of HD have been identified which delay or accelerate the onset HD or alter the severity of clinical phenotypes^14,18–21^. Modifiers identified from genome-wide association studies (GWAS) typically occur in genes encoding DNA mismatch repair proteins, with demonstrated effects on somatic *HTT* CAG expansion in mouse models of the disease^22–25^. Mismatch repair modifiers are hypothesized to alter the presentation of HD through modified excision-repair activity of the MutSβ complex, leading to altered somatic expansion in affected neurons of the brain^20,26,27^. However, phenotypic effects of mismatch repair modifiers are modest in HD clinical cohorts, and the causative biology of each modifier remains to be fully elucidated within the context of the human disease.

Modifiers within exon 1 of *HTT* accelerate the onset and progression of HD to a greater magnitude than mismatch repair modifiers. In canonical HD-causing alleles, the expanded CAG repeat in exon 1 is followed by two additional glutamine-encoding CAACAG codons, two proline-encoding CCGCCA codons, and a variable-length CCG repeat. In a subset of HD alleles, the interrupting CAACAG and CCGCCA codons are absent from this canonical sequence, resulting in uninterrupted CAG and CCG repeats in tandem^28^. Our group and others have shown that this CAG and CCG loss-of-interruption (CAG-CCG LOI) variant hastens the onset of HD motor symptoms by up to 12.5 years and accelerates clinical measures of disease progression when compared to patients with the canonical sequence^14,16,18,19,21,29^. The CAG-CCG LOI variant has also been shown to hasten striatal atrophy in premanifest individuals many years from predicted onset, suggesting this modifier acts early in the selective degeneration of HD-affected regions of the brain^30^. Understanding how the CAG-CCG LOI modifier accelerates the pathogenesis of HD in the striatum may therefore clarify causative mechanisms leading to HD^20,31^.

Direct correlations of somatic CAG expansion with phenotypic measures of HD remain poorly resolved. After accounting for age and inherited CAG repeat length, residual somatic expansion in blood of HD patients is reported to correlate with onset of HD^16^, but lacks clear association with striatal volume loss and CSF biomarkers NfL and PENK^30^. Association studies of somatic expansion in blood of HD patients have found many of the same mismatch repair loci that hasten or delay the onset of HD, yet the effects of these loci on blood somatic expansion are often inconsistent with their clinical effects^14^. Taken together, these data suggest that somatic expansion in blood may not accurately represent the dynamics of somatic expansion in HD-affected regions and neurons of the brain.

Medium spiny neurons (MSNs) of the striatum are selectively enriched for somatic CAG expansions and show higher expression of MutSβ subunits MSH2 and MSH3 relative to other striatal cell types^12^. Direct examination of somatic expansions in affected neurons of the brain may therefore be necessary to investigate the causality of somatic expansion on HD phenotypes and to determine whether genetic modifiers of HD act on somatic expansion in a cell type-specific manner. Striatal cell types exhibit widespread transcriptional changes in HD, but additional transcriptional changes observed in MSNs with more than 150 CAG repeats are hypothesized to directly lead to neuron dysfunction and loss^13^. The frequency of such highly expanded CAG repeats has not been reported in MSNs from HD patients, nor investigated in relation to genetic modifiers or phenotypes of HD.

Here, we apply complementary approaches to assess somatic *HTT* CAG expansion from peripheral blood, postmortem brain tissues, and isolated MSNs of HD patients with and without the CAG-CCG LOI modifier variant, and quantify MSN loss in CAG-CCG LOI donor caudate. While the CAG-CCG LOI variant does not increase the frequency of small somatic CAG expansions detectable in blood or bulk brain tissue, the modifier is strongly associated with increased proportions of MSNs with large somatic expansions and accelerated MSN loss. These results suggest that genetic changes in the *HTT* CAG repeat determine the rate of large somatic expansions in vulnerable human brain cell types and act to modify HD pathogenesis.

## Results

Mismatch repair modifiers of HD are hypothesized to act through altered somatic expansion of the *HTT* CAG repeat, but the causative mechanism of the CAG-CCG LOI variant in accelerating HD onset and progression is unknown. We therefore evaluated the impact of the CAG-CCG LOI variant on somatic expansion of the *HTT* CAG repeat in HD blood and brain tissue, as a potential biological explanation for accelerated HD pathology in these patients (**Figure 1**).

**Figure 1.**
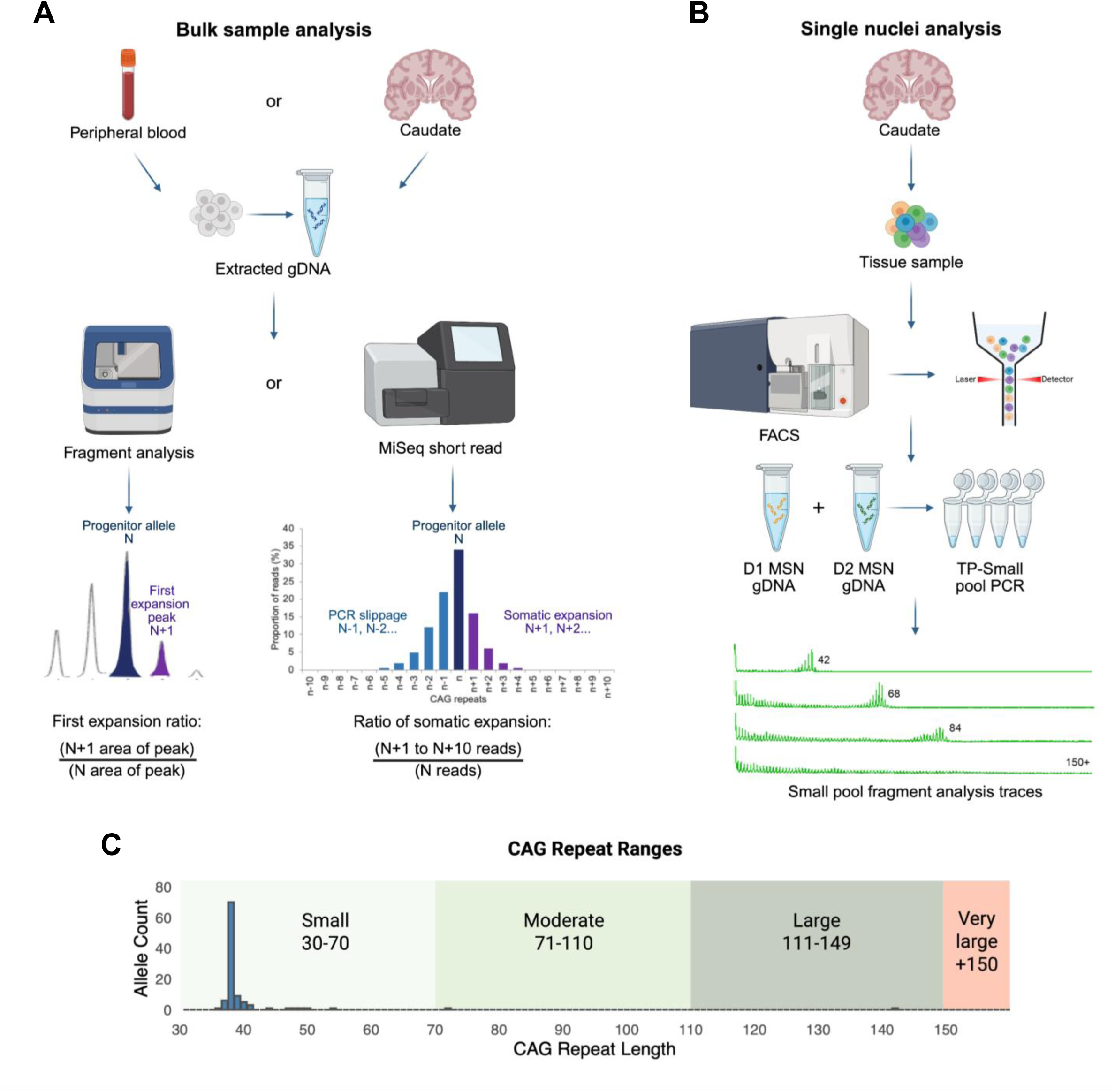
Methods for assessment of somatic *HTT* CAG expansion in HD tissue and sorted medium spiny neuron (MSN) nuclei. (**A**) Somatic expansion in bulk peripheral blood and postmortem brain tissue were assessed using fragment analysis to quantify the first expansion ratio (FER; expansions of +1 from inherited CAG repeat length) and MiSeq short-read sequencing to calculate the ratio of somatic expansion (RSE; expansions of +1 to +10 from inherited CAG repeat length). (**B**) To evaluate somatic expansion at the level of cell type, postmortem caudate tissue was sorted by Fluorescence-Activated Nuclear Sorting (FANS) using DRD1+ and DRD2+ RNA-complementary probes, followed by triplet-primed small-pool PCR (TPSP-PCR), enabling quantitative detection of somatic CAG repeat expansions across ranges. (**C**) The CAG repeat ranges include small (30–70), moderate (71–110), large (111–149), and very large (≥150) CAG expansions.

### Small somatic CAG expansions in blood do not correlate with age of onset and are reduced in HD patients with the CAG-CCG LOI variant

We first evaluated somatic expansion of the CAG repeat in peripheral blood, given the availability of this tissue from a large number of donors with known *HTT* CAG-CCG repeat sequence composition. For this analysis, we included 125 donors with the canonical sequence (uninterrupted CAG repeat range: 30-52) and 24 donors with the CAG-CCG LOI variant (uninterrupted CAG repeat range: 32-52). Among symptomatic donors included in the blood expansion analysis, the CAG-CCG LOI accelerated HD onset by 10.5 years relative to expected from uninterrupted CAG repeat length. We performed ultra-high depth MiSeq amplicon sequencing across the *HTT* CAG-CCG repeat region and assessed the ratio of amplicons with expansions of N+1 to N+10 to amplicons with the inherited CAG repeat length (N) in each donor (ratio of somatic expansion, RSE). For a larger set of 201 canonical donors and 37 CAG-CCG LOI variant donors, we also assessed the ratio of N+1 CAG expansion amplicons to inherited (N) CAG repeat amplicons from electrophoresis fragment peak heights (first expansion ratio, FER).

Small somatic expansions in the blood of HD patients evaluated by short read sequencing and by fragment length analysis increase with uninterrupted CAG repeat length and with age at sample collection (**Figure 2A**, **Supplemental Figure 1, Supplemental Figure 2A**). However, after controlling for uninterrupted CAG repeat length and age at collection, we detected no correlation between residual age of disease onset and residual somatic expansion in blood DNA of canonical donors or CAG-CCG LOI donors by either RSE or FER (**Figure 2B, Supplemental Figure 2B**). Surprisingly, residual somatic expansion was reduced in blood DNA from CAG-CCG LOI donors compared to canonical donors when adjusted for inherited CAG repeat length and age (*p=* 3.8×10^-4^, **Figure 2C**). When genotype was added to the model, the marginal mean of somatic expansion was reduced in blood of patients with the CAG-CCG LOI compared to the canonical sequence (*p*=6.4×10-4, **Figure 2D, Supplemental Table 1, model 1**). Similar findings were obtained from the electrophoresis-bases analysis, though reduced somatic expansion in CAG-CCG LOI variant alleles was not detectable by measurement of first expansion ratio (Supplemental Figure 2B-D, **Supplemental Table 2, model 1**).

**Figure 2.**
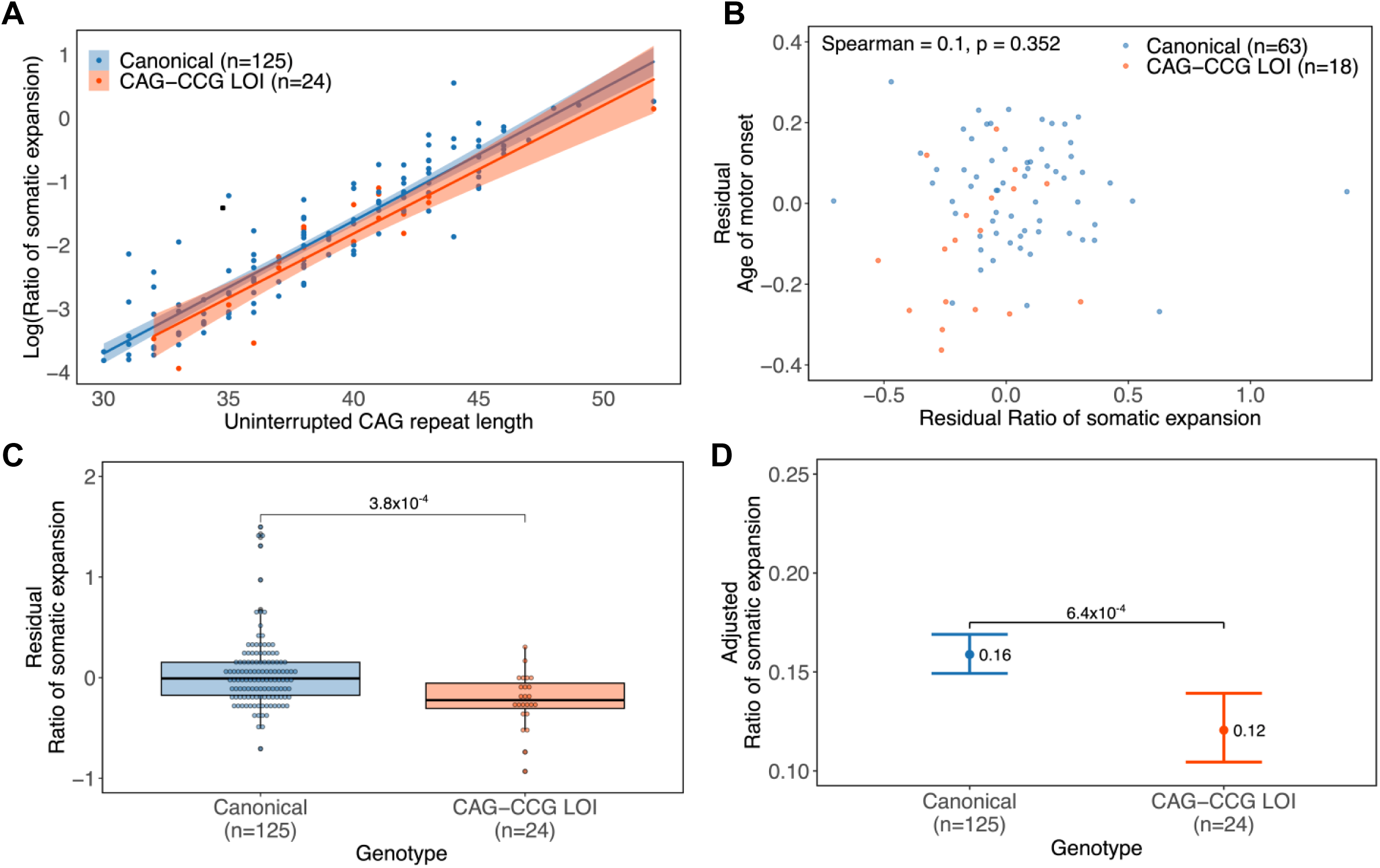
Small somatic CAG expansions are reduced in the peripheral blood of HD patients with the CAG-CCG LOI variant and do not correlate with age of onset. MiSeq-based bulk somatic instability in peripheral blood was examined in relation to genotype and age at motor onset. (**A**) Scatter plot of the uncorrected log-transformed ratio of somatic expansion versus uninterrupted CAG repeat length in peripheral blood DNA from 125 canonical and 24 CAG-CCG LOI variant donors. Shaded regions represent 95% confidence intervals. (**B**) Residual ratio of somatic expansion from 63 canonical and 19 CAG-CCG LOI variant donors with known motor onset, plotted against residual age at motor onset after correcting for uninterrupted CAG length, revealing no association between residual somatic expansion and residual motor onset (Spearman p-value: 0.352). (**C**) Residual ratio of somatic expansion for 125 canonical and 24 CAG-CCG LOI variant donors after adjusting for uninterrupted CAG length, age at sample collection, and their interaction. Residual somatic expansion was significantly lower in the CAG-CCG LOI variant donors (p-value: 3.8×10^-4^). (**D**) Estimated marginal means of the ratio of somatic expansion of the linear model including genotype, uninterrupted CAG repeat length, age at blood sample collection, and the CAG-age at sample collection interaction, showing a significantly reduced ratio of somatic expansion in the CAG-CCG LOI variant donors (p-value: 6.4×10^-4^). Error bars represent 95% confidence intervals.

### The CAG-CCG LOI variant does not increase the frequency of small somatic CAG expansions in bulk brain tissue of HD patients

Somatic instability of the *HTT* CAG repeat is tissue-specific, with higher rates of somatic expansion in affected brain regions relative to blood^8,15^. Equivocal effects of GWAS loci on HD onset and somatic expansion in blood suggest that blood DNA is a poor proxy for detection of disease-relevant somatic expansion in the brain^14^. To evaluate the effect of the CAG-CCG LOI modifier on somatic expansion in HD brain tissue, we compared caudate, putamen, frontal cortex, and cerebellum between 34 donors with the canonical sequence and 11 donors with the CAG-CCG LOI variant (uninterrupted CAG repeat range 38-44). Somatic expansion of the CAG repeat in brain tissues was assessed by short-read sequencing and fragment length analysis, consistent with the approaches used for blood. Among canonical donors, somatic expansion increases with uninterrupted CAG repeat length in all four regions by both RSE and FER, without significant relationship to duration of disease (**Figure 3A, 3C, 3E, 3G, Supplemental Tables 1 and 2, model 2**).

**Figure 3.**
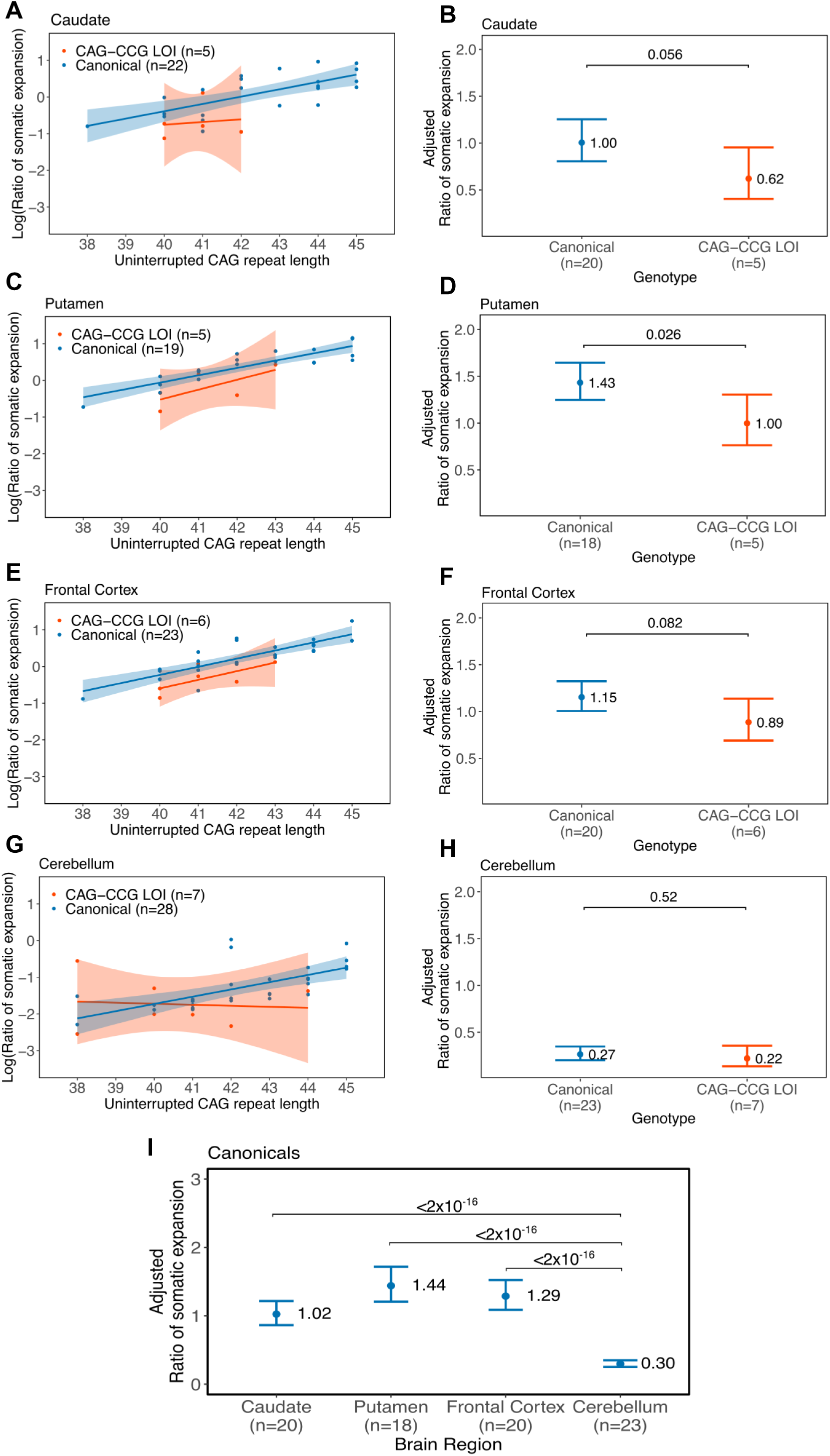
The CAG-CCG LOI variant does not increase small somatic expansions in brain tissue of HD patients. (**A, C, E, G**) The log-transformed ratio of somatic expansion varies by uninterrupted CAG repeat length in brain tissues from canonical and CAG-CCG LOI variant donors (**B, D, F, H**) Estimated marginal means of the ratio of somatic expansion, accounting for genotype, uninterrupted CAG repeat length, disease duration, and the interaction of CAG with disease duration. Significant differences in the adjusted ratio of somatic expansion were observed in the putamen and marginally in the caudate in donors with the CAG-CCG LOI (p-value: 0.026 and 0.056, respectively). Error bars represent 95% confidence intervals. (**I**) The ratio of somatic expansion, adjusted for uninterrupted CAG repeat length and age at donation, differs between caudate, putamen, and frontal cortex relative to the cerebellum in canonical donors (p<2×10^-16^).

Adjusted somatic expansion was calculated from linear models of RSE and FER in each brain tissue, controlling for uninterrupted CAG repeat length. Similarly to blood, Estimated marginal means of RSE were lower in CAG-CCG LOI putamen and trended towards reduction in the caudate by RSE, and estimated marginal means of FER were lower in caudate and trended towards reduction in the putamen by FER (**Figure 3B, 3D, Supplemental Figure 3B, 3D, Supplemental Tables 1 and 2, models 3-4**). No difference in adjusted somatic expansion was observed in frontal cortex or cerebellum of donors with the CAG-CCG LOI compared to donors with the canonical sequence, despite a positive correlation between CAG repeat length and somatic expansion across all examined brain tissues of both genotypes (**Figure 3E, 3G, Supplemental Figure 3E,3G, Supplemental Tables 1 and 2, models 5-6**). Estimated marginal means of somatic expansion were not different between genotypes in frontal cortex or cerebellum by either RSE or FER (**Figure 3F, 3H, Supplemental Figure 3F,3H**). Notably, adjusted somatic expansion of the CAG repeat is significantly lower in cerebellum compared to caudate, putamen, and cerebral cortex by both RSE and FER, as previously reported (**Figure 3I**, **Supplemental Figure 3I**)^8,15^.

### TPSP-PCR detects large somatic CAG expansions in HD caudate and enrichment in MSNs

PCR and sequencing of bulk tissues can result in poor detection of rare somatic variants, especially large CAG expansions present in HD brains^32,33^. We reasoned that our measures of small somatic expansion from bulk brain could obscure longer, less frequent somatic expansions that may be relevant to HD pathology in specific neurons and selectively impacted by the CAG-CCG LOI modifier. To investigate the presence of rare somatic expansions at the single-genome level, we developed a small-pool PCR assay to amplify and assess CAG repeat length from individual DNA molecules comprising *HTT* exon 1 sequence. This novel small pool assay incorporates triplet-priming of the CAG repeat, as commonly used in diagnostic ascertainment of expanded juvenile HD alleles, to detect highly expanded CAG repeat molecules that are not reliably amplified by conventional CAG repeat primer pairs^34,35^. When applied to DNA from bulk HD caudate, this novel triplet-primed small pool PCR (TPSP-PCR) approach yielded somatic distributions of CAG repeat molecules from unexpanded and expanded alleles in approximately equal proportions, indicating that amplification of genomic CAG repeat molecules is not biased by CAG repeat length in this assay (**Supplemental Figure 4A**). Among CAG repeat molecules from the expanded HD allele, somatic expansions could be measured across the range of CAG repeat lengths to ∼150 CAG, and detected beyond this threshold by stereotypical triplet-priming stutter (**Figure 4A-4B**, **Supplemental Figure 4B-5D**). We divided the range of somatically expanded CAG molecules into moderate (71-110), large (111-150 CAG) and very large (>150 CAG) somatic expansions^13,25^.

**Figure 4.**
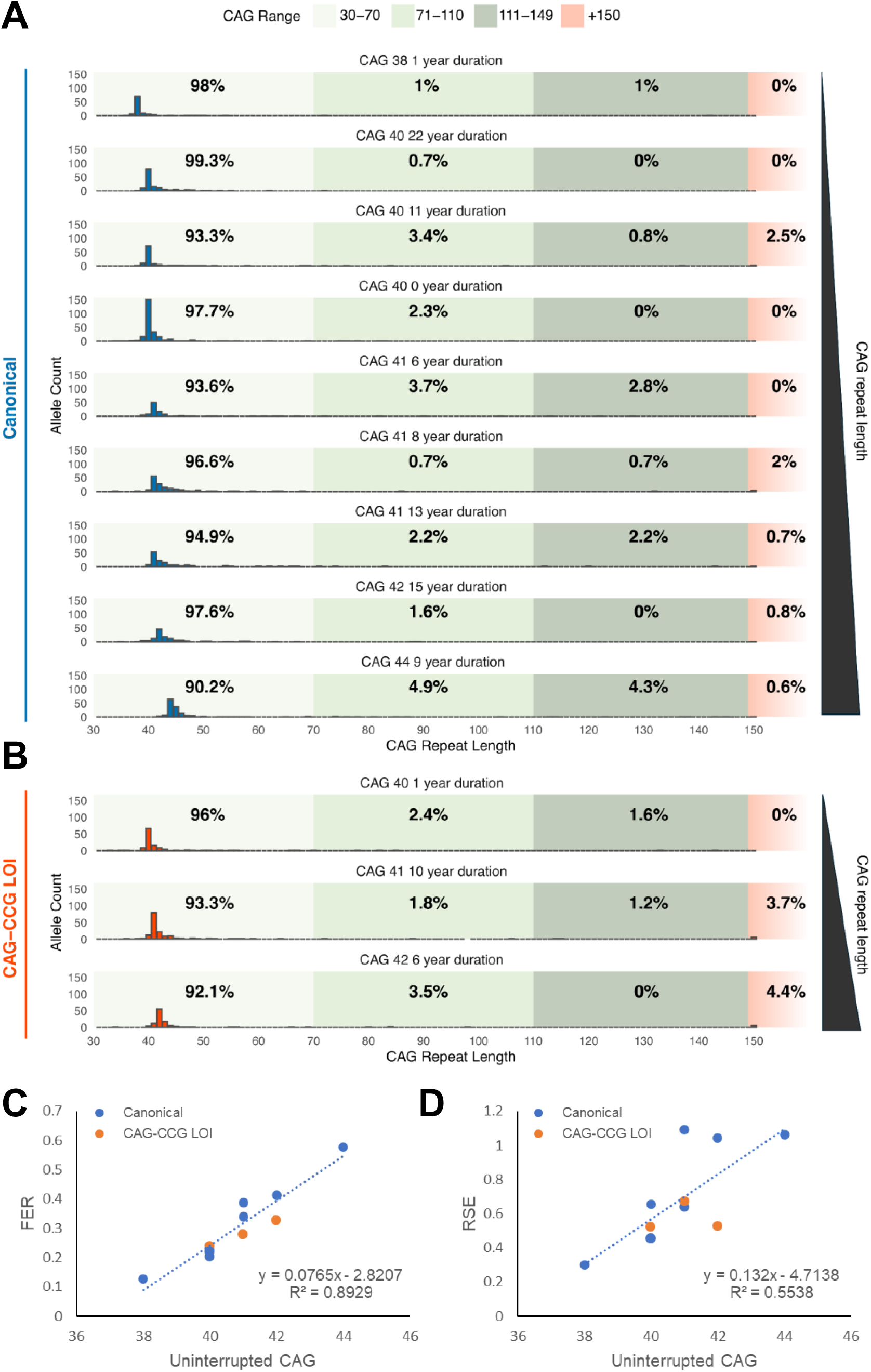
Triplet-primed small pool PCR (TPSP-PCR) shows no increase in small expansions in CAG-CCG LOI donors, but reveals large somatic CAG expansions in HD caudate. Small pool PCR detects moderate (70-110), large (111-150) and very large (>150) somatic CAG expansions at low frequency in bulk HD caudate from (**A**) canonical-sequence donors and (**B**) CAG-CCG LOI donors, ordered here by inherited CAG and by disease duration within each genotype group (canonical or CAG-CCG LOI. Linear modeling of somatic expansion in bulk caudate by TPSP-PCR showed significant increase with inherited uninterrupted CAG repeat length (p=0.0095), but no significant effect of CAG-CCG LOI genotype. In agreement with our short-read data, TPSP-PCR first expansion ratio (FER, **C**) and ratio of somatic expansion (RSE, **D**) showed no effect of CAG-CCG LOI genotype on small expansions in bulk HD caudate.

To compare somatic expansions in MSNs between donors with the CAG-CCG LOI and the canonical sequence, we selected caudate samples from donors of each genotype, closely matched for uninterrupted CAG repeat length and disease duration (**Supplemental Table 3**). TPSP-PCR analysis of bulk caudate from matched donors with the CAG-CCG LOI or canonical sequence confirmed that CAG-CCG LOI genotype has no effect on somatic expansion in the range of N+1 to N+10 CAG, despite increasing somatic expansion in this range with inherited CAG repeat length (**Figure 4C, D**). Linear modeling of somatic expansions detected in bulk caudate by TPSP-PCR showed a significant effect of inherited uninterrupted CAG repeat length, but no significant effect of CAG-CCG LOI genotype (**Supplemental Table 4, model 1)**.

Recent single-cell RNA-seq analyses of HD caudate show that at later disease stages, MSNs are infrequent among caudate cell types due to progressive MSN loss ^12,13^. Small-pool PCR distributions generated from bulk caudate may therefore overrepresent *HTT* copies from striatal cell types other than MSNs, and are not comparable across disease stages due to different cell type compositions. To determine the distribution of somatic CAG repeat lengths in MSN genomes, we employed FANS using RNA probes directed to *PPP1R1B*, encoding the MSN-specific marker DARPP-32^12,36^. We performed short-read sequencing from DNA of unsorted caudate nuclei and sorted MSN nuclei, and then compared to single-molecule TPSP-PCR CAG repeat distributions from the same samples. The somatic CAG distribution of unsorted caudate nuclei by TPSP-PCR resembled the somatic CAG distribution by short read sequencing, with added detection of very large somatic expansions >150 CAG and the elimination of smaller PCR slippage products from the expanded HD allele (**Figure 5A**). From sorted MSN nuclei, TPSP-PCR analysis showed a broad distribution of expanded CAG repeat lengths comparable to short-read sequencing but an unexpectedly high frequency (>20%) of somatic expansions to >150 CAGs which were undetectable with short-read sequencing (**Figure 5B**). TPSP-PCR thus enables detection of large somatic CAG expansions, including somatic expansions >150 CAG, that are highly enriched in caudate MSN nuclei relative to other abundant striatal cell types.

**Figure 5.**
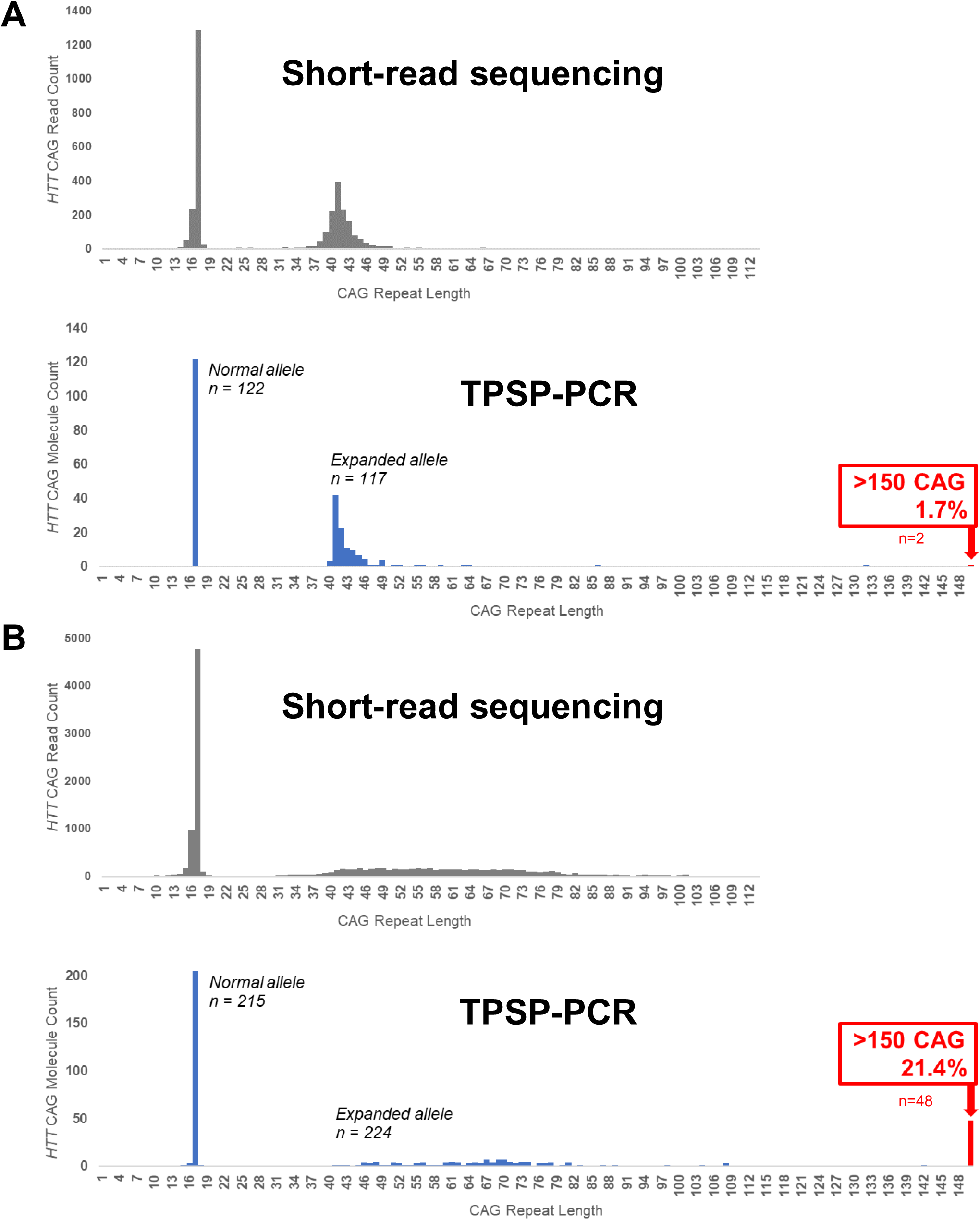
Comparison of short-read *HTT* CAG repeat amplicon distributions and TPSP-PCR single-molecule *HTT* CAG repeat distributions. (**A**) Comparison of short-read and TPSP-PCR CAG distributions from unsorted HD caudate nuclei and (**B**) from *PPP1R1B* (DARPP-32+) sorted caudate MSN nuclei.

### Large somatic expansions in MSNs increase in proportion with inherited CAG length, disease duration, and CAG-CCG LOI genotype

When applied to nuclei expressing *PPP1R1B* and *DRD1*, TPSP-PCR revealed that somatic distributions of large (111-150) and very large (>150) CAG expansions are variable among canonical D1 MSNs, increasing with duration of disease and inherited CAG repeat length (**Figure 6A**). The proportion of caudate MSNs with large and very large somatic expansions in a canonical donor 1 year after motor onset were 3.4% and 0.8%, respectively, increasing to 10.7% and 16.1% in a canonical donor 15 years after motor onset (Fisher’s exact test p=0.037 and p=1.36×10^-5^, respectively). The proportion of caudate MSNs with HD repeats ≤70 CAG correspondingly declined in canonical donors with disease duration, from 84.7% at 1 year to 39.3% at 15 years, as the proportion of large and very large expansions increased.

**Figure 6.**
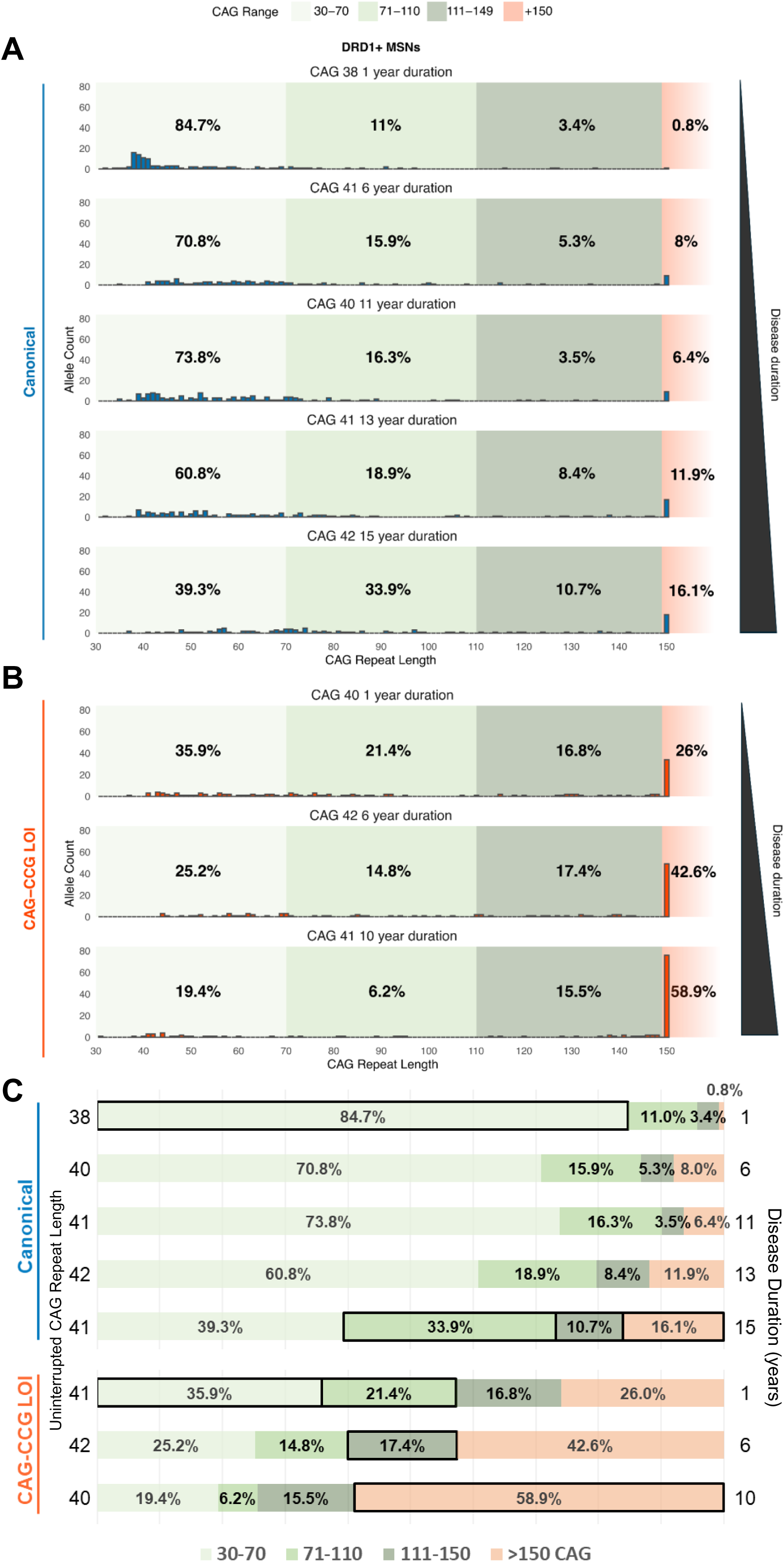
Large somatic expansions in MSNs increase in proportion with disease duration and CAG-CCG LOI genotype. Single-molecule *HTT* CAG repeat distributions in caudate D1 MSNs from (**A**) canonical-sequence HD donors and (**B**) CAG-CCG LOI variant HD donors. (**C**) Grouped proportions of small (<50), moderate (51-110), large (111-150), and very large (>150) somatic CAG expansions in D1 MSNs. Proportions boxed in black represent the highest proportion of a given CAG length group among canonical donors or among CAG-CCG LOI donors. Large (111-150) and very large (>150) somatic CAG expansions are enriched by duration between canonical donors 1 year and 15 years from motor onset (*p=*0.03744 for large expansions and *p*=1.365x10^-5^ for very large expansions). Large (111-150) and very large (>150) somatic CAG expansions are further enriched by CAG-CCG LOI variant status in donors with 1 year disease duration (*p*=6.35x10^-4^ large and *p*=9.133x10^-10^ very large expansions) and donors with 10 year disease duration (*p*=1.186x10-3 large and *p*<2.2x10^-16^ very large). In canonical CAG molecule distributions, somatic CAG repeat length is significantly increased by inherited uninterrupted CAG repeat length (p=0.0235), and by the interaction of inherited uninterrupted CAG repeat length with disease duration (p=0.0322). In linear mixed modeling of the combined data set, somatic CAG repeat length is strongly increased by CAG-CCG LOI status (p=2.18x10^-10^), as well as by the interaction of disease duration with CAG-CCG LOI status (p=0.0393).

Single-molecule CAG distributions of MSNs from donors with the CAG-CCG LOI were strikingly more expanded when compared to canonical donors (**Figure 6B**). Large (111-150) and very large (>150) somatic CAG expansions were enriched in MSNs by CAG-CCG LOI genotype between matched donors with 1 year disease duration (large frequency: 3.4% (n=4/118 canonical) vs 16.8% (n=22/131 CAG-CCG LOI), p=6.35×10^-4^ Fisher’s exact test, and very large frequency: 0.8% (n=1/118 canonical) vs 26.0% (n=34/131 CAG-CCG LOI), p=9.13×10^-10^ Fisher’s exact test); between matched donors with 6 year disease duration (large frequency: 5.3% (n=6/113) vs 17.4% (n=20/115), p=5.91×10^-3^, and very large frequency: 8.0% (n=9/113) vs 42.6% (n=49/115), p=9.50×10^-10^); and between matched donors with 10/11 year disease duration (large frequency: 3.5% (n=5/141) vs 15.5% (n=20/129) p=1.186×10^-3^, and very large frequency: 6.4% (n=9/141) vs 58.9% (n=76/129) p<2.2×10^-16^). Remarkably, the observed proportions of large and very large somatic expansions in CAG-CCG LOI caudate MSNs exceeded that of canonical caudate MSNs at all disease stages, and were increased ∼5-fold relative to canonical donors at matched disease duration and inherited CAG length (**Figure 6C**). As in canonical MSNs, the proportion of very large somatic expansions in CAG-CCG LOI MSNs increases with disease duration from 26% of MSNs at 1 year disease duration to 58.9% at 10 years. Thus, at 10 years from motor onset in a donor with inherited 41 CAG and the CAG-CCG LOI variant, a majority of caudate MSNs have somatic expansions greater than 150 CAG repeats. We observed no difference in the patterns of instability between D1 and D2 MSN subgroups in either canonical or CAG-CCG LOI donors, suggesting that MSNs of the direct and indirect pathway have similar trajectories of somatic expansion (D1 vs D2 CAG distributions nonsignificant in each donor by Kolmogorov-Smirnov Test, **Supplemental Figure 5**).

To better understand the effect of different variables associated with increased somatic expansion in MSNs, we performed linear modeling of CAG molecule distributions from canonical donors and also the combined data set including CAG-CCG LOI donors (**Supplemental Table 4, models 1 and 2**). Across canonical CAG molecule distributions, somatic CAG repeat length was significantly increased by inherited uninterrupted CAG repeat length (p=0.024), and the interaction of disease duration with inherited uninterrupted CAG repeat length (p=0.032). In the combined data set, somatic CAG repeat length was marginally significantly increased by disease duration (p=0.051) as well as the interaction of disease duration with inherited CAG repeat length (p=0.0393), but was very strongly increased by CAG-CCG LOI genotype (p=2.18×10^-10^) and the interaction of disease duration with CAG-CCG LOI genotype (p=0.016). These results suggest that the CAG-CCG LOI exerts the strongest effect on somatic expansion in MSNs among all tested variables.

### The CAG-CCG LOI is associated with earlier MSN loss and earlier DRD2+ MSN depletion

Large (111-150 CAG) and very large (>150 CAG) somatic expansions have been associated with widespread transcriptional changes and are hypothesized to directly cause MSN loss^13,25^. We therefore sought to quantify the absolute abundance of DARPP-32+ MSNs in fixed caudate sections from a larger cohort of CAG-CCG LOI and canonical HD donors, including donors analyzed in our FANS and TPSP-PCR experiments (**Figure 7A**, **Supplemental Table 3**).

**Figure 7.**
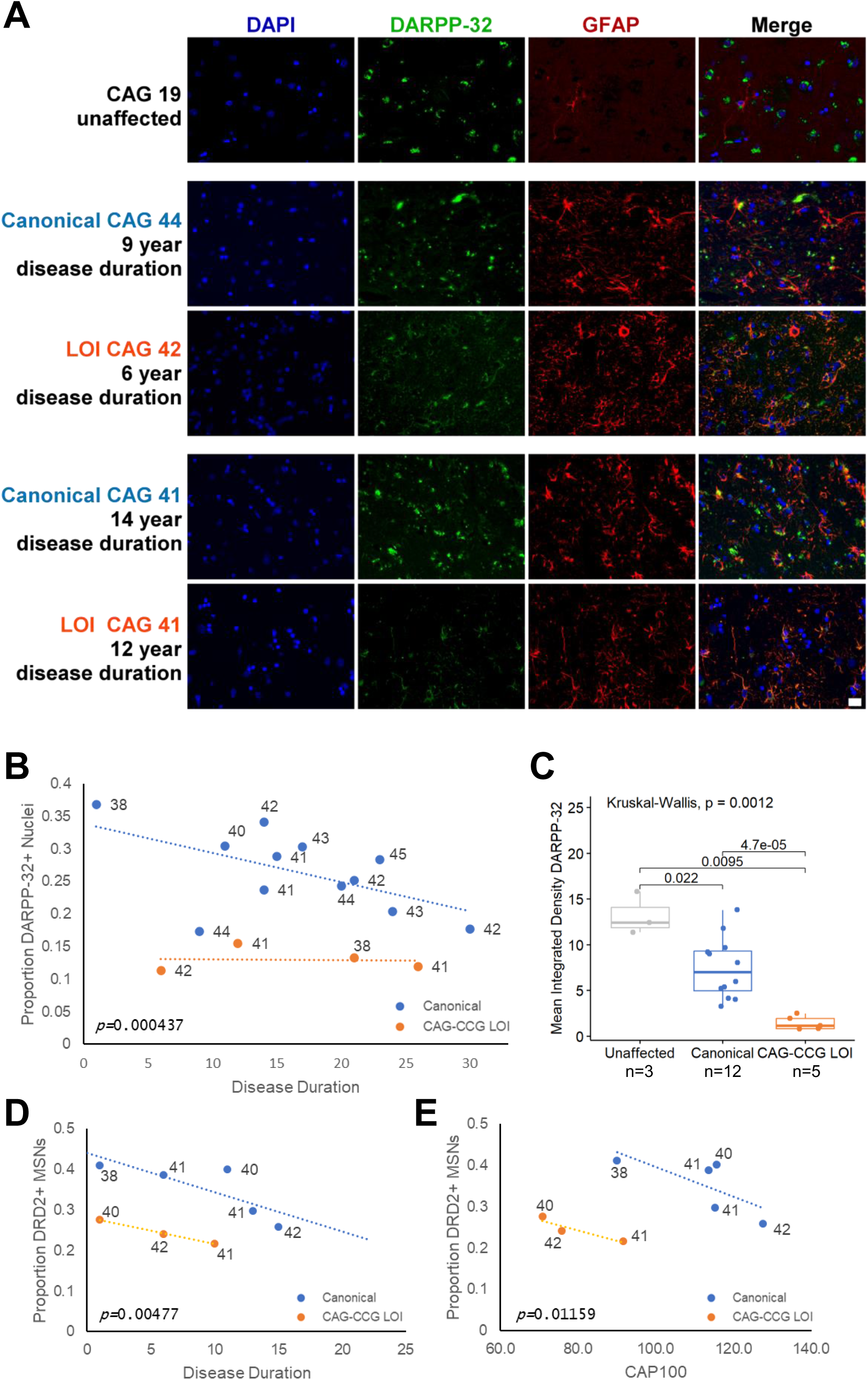
The CAG-CCG LOI is associated earlier MSN loss and earlier D2 MSN depletion. (**A**) IF staining of DAPI, DARPP-32 and GFAP in HD caudate sections from canonical-sequence and CAG-CCG LOI donors. (**B**) Proportion of DARPP-32+ nuclei among all counted caudate nuclei is reduced in CAG-CCG LOI donors by linear regression (p=4.4x10^-4^). Proportion of DARPP-32+ nuclei in unaffected controls was 44% (not shown). (**C**) Mean integrated DARPP-32 intensity by donor genotype. DARPP-32 intensity per cell is lower in HD-affected canonical-sequence caudate (n=12) than unaffected controls (n=3, p=0.022, t-test). DARPP-32 intensity per cell is lower in HD-affected CAG-CCG LOI caudate (n=5) than in affected canonical sequence caudate (n=12, p=4.7x10^-5^, t-test). (**D**) Proportion DRD2+ MSNs among all MSNs by disease duration is reduced in CAG-CCG LOI donors relative to canonical by linear regression (p=4.8x10^-3^). (**E**) Proportion DRD2+ MSNs among all MSNs by CAP100 score is reduced in CAG-CCG LOI donors relative to canonical by linear regression (p=0.012), but earlier age of death in CAG-CCG LOI donors with earlier onset precludes direct comparison by CAP100.

DARPP-32+ MSN counts in caudate from canonical donors declined as a proportion of all nuclei with disease duration, from ∼35% of caudate nuclei at onset to ∼20% of caudate nuclei in later disease stages. MSN counts in caudate sections from CAG-CCG LOI donors were significantly lower than counts from canonical donors across all time points (p=4.37×10^-4^, **Figure 7B**) and DARPP-32+ signal was reduced among positive nuclei (p=4.7×10^-5^, **Figure 7C**). Sorting of DARPP-32+ DRD1+ nuclei and DARPP-32+ DRD2+ nuclei from HD caudate allowed comparison of the proportion of DRD2+ to DRD1+ MSNs in these donors. The relative proportion of DRD2+ MSNs in canonical donors declined with disease duration, from ∼40% of caudate MSNs in early HD to <30% at advanced duration, reflecting the increased vulnerability of D2 MSNs relative to D1 MSNs over the course of the disease (**Figure 7D**). Strikingly, the proportion of DRD2+ MSNs was lower in CAG-CCG LOI caudate than canonical caudate when compared by disease duration (p=4.77×10^-3^), suggesting greater striatal D2 MSN loss than D1 MSN loss in donors with the CAG-CCG LOI variant, despite no detectable difference in somatic expansion between these MSN subpopulations. Comparison of caudate MSN counts between CAG-CCG LOI donors and canonical donors by CAP100 was uninformative due to earlier death of CAG-CCG LOI donors at similar inherited CAG lengths, reflecting the dramatically earlier HD onset in donors with the CAG-CCG LOI variant (**Figure 7E**).

## Discussion

People who develop HD are born with 36 or more uninterrupted *HTT* CAG repeats, and are usually asymptomatic in the first decades of life. The pathology and clinical features of HD emerge gradually, in parallel with the slow and persistent loss of striatal MSNs. The CAG-CCG LOI is the strongest known genetic modifier of HD, accelerating onset of the disease by more than 12 years, hastening progression in motor impairment, and worsening striatal volume loss^30^. Our interrogations of somatic expansion in blood, brain, and striatal MSNs of donors with and without the CAG-CCG LOI show this modifier does not increase small expansions in blood or bulk brain tissues as previously reported^14,37^, yet profoundly increases the proportion of genomic large (111-150) and very large (>150) CAG expansions in affected striatal MSNs. These findings suggest that large somatic CAG repeat expansions in MSNs, which cannot be detected and appropriately quantified in bulk striatal tissue, contribute to HD striatal pathology and may mediate the disease-hastening effect of the CAG-CCG LOI modifier.

It has been proposed that MSNs with >150 CAG are uniquely vulnerable and rapidly cause MSN pathology and loss^13^. Our study provides direct evidence that somatic expansions >150 are rare in MSN genomes in the early clinical stages of HD, yet accumulate with time from onset and in greater magnitude with the CAG-CCG LOI modifier. Expansions >150 CAG are observed in <1% of MSNs from a canonical donor one year after onset, despite classical presentation of HD and notable MSN loss (37% MSNs among all caudate cells versus 44% in controls), suggesting that expansions >150 CAG do not uniquely drive MSN dysfunction and loss in early HD. In contrast, a CAG-CCG LOI donor only one year after onset exhibits a 5-fold greater proportion of large (111-150) and very large (>150) CAG expansions in MSNs, and this enrichment increases as disease duration progresses. Taken together, these data suggest that the overall proportion of large genomic CAG expansions in MSNs may reflect the rate of HD pathology, but that susceptibility of individual MSNs to dysfunction and loss can occur at somatic lengths below 150 CAG.

Is there a specific cytotoxic threshold of somatic expansion that causes MSNs to die? Repeat lengths of MSNs that have already been lost are not observed in our study. Very large expansions >150 CAG are rare in early-stage canonical MSNs, however these very large expansions accumulate in MSN genomes with longer duration of disease and with the CAG-CCG LOI modifier, suggesting that MSNs with >150 CAG are not rapidly lost and can persist in the HD brain. Additionally, we observe no significant difference in large (111-150 CAG) and very large (>150 CAG) somatic expansions between D1 and D2 MSNs, despite greater vulnerability of D2 MSNs in HD and accelerated D2 MSN loss in CAG-CCG LOI donors, suggesting that the proportion of somatic expansions >150 is not the sole determinant of MSN loss. Our data thus support a model in which large somatic expansions increase MSN susceptibility to dysfunction and loss, but that specific CAG lengths causing MSN dysfunction include repeats below 150 CAG. Our data are also compatible with a model in which a subset of MSNs with repeats beyond 150 CAG are comparatively spared, leading to their accumulation among MSNs with advancing disease and the CAG-CCG LOI genotype. Although TPSP-PCR enables the detection of very large genomic CAG expansions and quantification of their total frequency among MSNs, individual repeat lengths beyond 150 CAG and their relative representation among surviving neurons could not be resolved.

Extreme constitutive lengths of the *HTT* CAG repeat are reported to reduce expression of the corresponding mRNA transcript and protein in cell and mouse models of HD^38,39^. Inferences of genomic *HTT* CAG repeat frequency through RNA-seq approaches may therefore underrepresent very large genomic CAG expansions (>150 repeats), particularly in MSN populations from HD patients where such somatic expansions are common. Our data suggests that MSNs with very large genomic CAG expansions persist within a smaller and smaller pool of surviving caudate neurons and may elude RNA-based detection.

The absence of detectable differences in small somatic instability in blood from CAG-CCG LOI donors versus canonical donors, despite the profound effect of the CAG-CCG LOI on MSN somatic expansion, suggests that assessment of blood DNA is of limited use as a biomarker for disease-relevant expansion. We additionally detect no correlation between residual CAG expansion and disease onset in bulk peripheral blood mononuclear cells (PBMCs), arguing that somatic expansion in blood is not an accurate correlate of disease onset.

Nuclei sorting methods employed in our study rely on expression of target RNA. *PPP1R1B* expression has been shown to be reduced in MSNs with >150 CAG^13^, which could lead to undercounting of MSNs with >150 repeats in our small pool experiments. However, this would not change our conclusion that large expansions increase in proportion with disease duration and CAG-CCG LOI genotype. In direct DARPP-32+ MSN count experiments, reduced *PPP1R1B* expression could also result in undercounting MSNs without DARPP-32. However, the CAG-CCG LOI is still associated with earlier loss of DARPP-32+ MSNs and reduced proportion of DARPP-32+ DRD2+ MSNs. Technical limitations that may be introduced by reduced or lost MSN identity markers underscore the need for *in situ* spatial transcriptomics with redundant markers to confirm cell type identity concurrent with somatic CAG expansion length.

MSN loss is reported to be variable among controls and HD donors with CAP scores near onset^13^. Our observation that MSN counts are lower in HD patients with the CAG-CCG LOI suggests that the onset and progression of HD may be a consequence of sustained MSN dysfunction at various somatic CAG repeat lengths, rather than percent MSNs lost beyond a specific toxic CAG threshold. It is noteworthy that somatic expansion and early DARPP-32+ MSN loss are not prominent features of many HD mouse models^40,41^. In HD mouse models without notable somatic expansion or MSN loss^40,42^, the constitutive expanded CAG repeat length could result in MSN pathology corresponding to a narrow range of MSNs found in HD patients at a given moment of the disease. Appropriate *in vivo* models of HD must include somatic expansion to accurately recapitulate what is observed in HD patients, especially for study of the CAG-CCG LOI and its phenotypic effects.

Our data do not exclude the possibility that the modifier effect associated with the CAG-CCG LOI is mediated, at least in part, through mechanisms other than increased somatic expansion. Mechanisms compatible with synonymous mutations of the CAG repeat sequence include direct RNA toxicity, RAN translation, ribosome stalling, and other possibilities mediated by changes to DNA or RNA secondary structure. Mechanisms of other known LOI modifier variants, including synonymous loss of interruption to the CAG repeat (CAG LOI) or the CCG repeat (CCG LOI) independently, have yet to be established. Future experiments must evaluate whether other loss of interruption modifier variants also result in increased somatic expansion in MSNs of HD patients or appropriate human-derived cell lines^43^.

In conclusion, we show that the HD genetic modifier with the largest magnitude of effect, the CAG-CCG LOI, is associated with an increased proportion of large somatic expansions in HD striatal MSNs, lower MSN counts, and earlier loss of D2 MSNs of the indirect pathway. Peripheral blood DNA does not capture increased somatic expansion in MSNs, suggesting that blood DNA is a poor biomarker for disease-relevant somatic expansion in affected neurons. While the underlying mechanism by which the CAG-CCG LOI modifier influences somatic expansion remains unresolved, our results suggest HD pathogenesis can be modified through introducing local genetic changes to the *HTT* CAG repeat region that alter its expansion in vulnerable cell types in human brain. Our study further underscores the need for cell type-specific studies across other repeat expansion disorders that preferentially affect specific neuron populations.

## Acknowledgements

We wish to acknowledge support of the Huntington Disease Foundation, the Huntington Disease Society of America, and the Huntington Society of Canada for funding this research. We also recognize the contributions of the UBC HD Biobank and the NZ Neurological Foundation Human Brain Bank for providing human tissue for this study. This work is not possible without the generous brain donations of HD patients and their families. We also thank Dr. Nicholas Caron and Dr. Christopher Pearson for helpful conversations pertaining to this work.

## Methods

### Donor Recruitment and Tissue Selection

To examine somatic expansion in blood, we identified a cohort of donors from the UBC HD Biobank, Ruhr University Bochum, and the University of Auckland with known CAG-CCG LOI variant genotype and available blood DNA (n = 37), including both HD-affected and unaffected donors. We assembled a comparison cohort of individuals with canonical alleles based on prior screening (n = 201), including both HD-affected and unaffected donors. The comparison group included the same range of uninterrupted CAG repeat lengths as the CAG-CCG LOI cohort (range 32-52 for both groups, mean CAG repeat length 40.1±4.3 for canonical HD, 38.6±4.1 for CAG-CCG LOI), and mean age at blood sample collection was well matched across donors at at each CAG repeat length.

To examine somatic expansion in brain tissue, we identified 11 brain donors with the CAG-CCG LOI variant from brain banks at the University of British Columbia and the University of Auckland. We then assembled a cohort of 35 canonical HD brain donors from the UBC HD Biobank at the University of British Columbia which were matched as closely as possible to the CAG-CCG LOI donors for uninterrupted CAG repeat length, disease stage, and disease duration. Vonsattel staging was available for most brains, and where unavailable was noted with approximate disease stage (early, mid, or late) based on the observed degree of caudate atrophy. DNA was extracted from frozen tissue sampled from the cerebellum, caudate, putamen, and frontal cortex of each donor as available.

To study single-molecule somatic expansion, a subset of HD brain donors with frozen caudate tissue was selected from the above cohort with best available matching by inherited uninterrupted CAG repeat length, disease duration, and neuropathological grade, comprising 7 canonical donors and 4 CAG-CCG LOI donors (Supplemental Table 1). Suitable caudate samples were then processed for fluorescence-activated nuclei sorting (FANS) as described below. Samples from 2 canonical donors and 1 CAG-CCG LOI donor subsequently failed quality control for FANS and were excluded.

For characterization of medium spiny neuron counts, a companion set of formalin-fixed paraffin-embedded (FFPE) caudate tissue sections was assembled from brain banks at the University of British Columbia and the University of Auckland, comprising 3 unaffected control donors, 12 canonical HD donors, and 4 CAG-CCG LOI donors, with partial overlap to the nuclei-sorted frozen caudate donor set.

### Capillary Electrophoresis and MiSeq Assessments of Somatic Expansion

For fragment length analysis of somatic CAG expansion, we amplified the *HTT* CAG repeat region from 60ng of input DNA using a fluorescently-labelled primer pair and reaction conditions typically used for CAG repeat genotyping (forward primer: HD344F-HEX, 5’-HEX-CCTTCGAGTCCCTCAAGTCCTTC; reverse primer: HD450R-PT, 5’-GTTTGGCGGCGGTGGCGGCTGTTG) (Andrew *et al.* 1994a, Semaka *et al.* 2013a). We performed all reactions in triplicate, diluted samples 1:60 prior to capillary electrophoresis using the ABI Prism 3500xl Genetic Analyzer, and analyzed results using GeneMapper5 software. We measured somatic expansion from blood and brain DNA samples using first expansion ratio (FER), a comparison of the peak area of the progenitor allele to the area of the peak representing an expansion of 1 CAG repeat.

The *HTT* short-read sequencing was performed using a previously established ultra-deep high-throughput protocol developed to precisely characterize the *HTT* exon 1 repeat tract (Ciosi et al. 2018). For the library preparation, 20ng of genomic DNA from both the peripheral blood and brain tissue samples was used, following the established protocol (Ciosi et al. 2018). Sequencing was conducted on a Illumina MiSeq platform.

Genotyping was carried out using a custom Galaxy pipeline. FASTQ files were demultiplexed with Cutadapt (maximum error rate 0.1) to isolate *HTT* repeat tract reads, followed by adapter trimming (maximum error rate 0.39). Trimmed reads were aligned using BWA-MEM (gap extension penalty 4.4) to synthetic *HTT* reference sequences generated using RefGeneratr, comprising 4,000 *HTT* references spanning 1–200 CAG and 1–20 CCG repeats and known sequence variants. Alignments were output in BAM format, converted to SAM using SAMtools and filtered to retain uniquely mapped reads (MAPQ > 0). Quality control was performed after each step using FastQC, assessing sequence quality, GC content, duplication levels, and overrepresented sequences to ensure data integrity. Filtered reads were then visualized using Tablet to confirm CAG repeat size and determination allele sequence structure of inherited expanded and unexpanded alleles.

For assessments of somatic CAG expansion, the ratio of CAG repeat somatic expansions of HD expanded alleles was determined from the MiSeq read count distributions as described previously (Ciosi *et al*. 2019). Ratio of somatic expansion (RSE) was calculated for each sample as the ratio of the sum of N+1 to N+10 reads to reads comprising the inherited uninterrupted CAG repeat length (N).

### Triplet-Primed Small Pool PCR Assessments

Triplet-primed small pool PCR (TPSP-PCR) was used to amplify CAG repeats from single-molecule *HTT* DNA inputs irrespective of CAG repeat length. DNA extracted from sorted and unsorted nuclei were serially diluted with Ultrapure distilled water (Invitrogen) and then added to a small pool master mix such that each 5uL reaction in each well contained between 3.2-6.4 pg of input DNA, corresponding to 0.5-1 genome equivalents per reaction. Each reaction plate included 12 randomly distributed negative controls, as well as a ladder of source DNA with known concentrations as positive controls. Following PCR, capillary electrophoresis was performed with the genetic analyzer ABI 3500xl Bioanalyzer, using 1.5uL-2.0uL of undiluted product from each reaction well mixed with 9uL of formamide.

GeneMapper6 software was used to analyze capillary electropherograms from each small-pool reaction. Wells without *HTT* locus-specific amplification were counted and compared to the expected number of empty reactions based on input DNA using a Poisson distribution. For each triplet-primed CAG repeat molecule amplification, the highest peak among the largest cluster of products showing contiguous stutter to the minimum length of 5 CAG was selected as the genotype. Where no apparent cluster of peaks corresponding to the genotype was evident but triplet-primed stutter was present up to 150 CAG and beyond, the genotype was assigned as >150 CAG for analytical purposes.

To exclude sequence bias in our initial triplet-primed primer set, an alternative triplet-primed primer set was designed to encompass both the CAG and CCG repeats, with triplet-priming instead directed to the 5’ end of the *HTT* CAG coding sequence. TPSP-PCR runs with this alternative primer set showed identical length distributions of somatically expanded CAG molecules as determined with the initial primer set, including identical frequencies of very large (>150 CAG) expansions observed in MSN DNA from donors with the canonical sequence or the CAG-CCG LOI sequence. These experiments with an alternative triplet-primed primer pair demonstrate that detection of highly expanded CAG repeats in our study are not primer-pair dependent.

### Fluorescence-Activated Nuclei Sorting (FANS) of Medium Spiny Neurons

Nuclei were isolated from 75-150mg of caudate from each donor as described previously (see Matlik *et al*. 2024, *Nature Genetics* and Pressl *et al*. 2024 *Current Protocols*). Briefly, cell nuclei were separated from homogenized caudate tissue by density gradient ultracentrifugation.

Isolated nuclei were resuspended in homogenization buffer (150 mM KCl, 5 mM MgCl2, 20 mM Tricene-KOH pH 7.8, and 250 mM sucrose with protease and RNase inhibitors) and fixed with 1% formaldehyde for 8 min at room temperature followed by quenching with 0.125 M glycine for 5 min. The nuclei were centrifuged 5 min at 1000 x *g* and the pellet washed with homogenization buffer, then centrifuged again 5 min at 1000 x *g* and the pellet washed in permeabilization buffer (1X PBS, 0.05% TritonX-100, 0.5% BSA, and RNase inhibitors). The nuclei were centrifuged a final 5 min at 800 x *g* and the pellet washed in permeabilization buffer, followed by incubation for 30 min at room temperature on a shaker.

For labeling neuronal nuclei, the PrimeFlow labeling kit (ThermoFisher, #88-18005-210) was used according to manufacturer’s instructions but with RNase inhibitors included in each incubation step. Probes specific to DRD1 (Alexa Fluor 647, ThermoFisher #VA1-3002351-PF), DRD2 (Alexa Fluor 488, ThermoFisher #VA4-3083767-PF) and PPP1R1B (Alexa Fluor 568, #VA10-3266354-PF) were used to label D1 MSN (647+, 568+, 488−, large) and D2 MSN nuclei (647−, 568+, 488+, large) populations. For sorting, PrimeFlow-labled nuclei were resuspended in DAPI buffer (1X PBS, 0.5% BSA, 0.5 μg/mL DAPI) and separated with SONY MA900 Cell Sorter (software ver. 3.0.5). Aggregates of nuclei were excluded based on higher intensity of DAPI staining.

DNA was extracted from sorted nuclei using the Qiagen Allprep DNA/RNA FFPE kit following manufacturer’s instructions, and quantified by Qubit fluorometer. Typical DNA yield was 1-2pg per counted nucleus.

### Immunofluorescence Staining and Nuclei Counting of Striatal Tissues

FFPE slides prepared from donor caudate were deparaffinized in xylene, followed by decreasing ethanol baths. Antigen retrieval was performed by Proteinase K digestion (1ug/ml in 1mM CaCl2/50mM Tris-HCl buffer (pH 7.6) for 40mins @ 37C) followed by boiling in citrate buffer (10mM Sodium Citrate, pH 6.0) for 15 mins in a pressure cooker, and aggregate unmasking done by formic acid (98%) treatment for 7 min.

Treated slides were blocked using ASE blocking buffer (30mins at room temperature) and labeled with primary Rat-DARPP32 (Novus, MAB4230, 1:200) in ASE incubation buffer (overnight at 4C, see Rosas-Arellano et al. 2016, *Histochem Cell Biol*). Primary labeled slides were washed in TBS-T and then incubated with secondary Goat-anti-rat 488 (Thermo, A11006, 1:500) and conjugated primary Mouse-anti-GFAP-Cy3 (Sigma, C9205, 1:1000) (2 hours at room temperature). Slides were counterstained with DAPI, and treated with Autofluorescence Quencher (Vector, SP-8400-15). Slides were mounted with Prolong Gold antifade mounting media (Thermo, P36930).

Imaging work was supported by the Imaging Core (RRID:SCR_026573) at BC Children’s Hospital Research Institute (BCCHR). Images were taken on the Olympus BX61 wide field fluorescence microscope at 20x magnification. Resulting images were processed using Cell Profiler (see Stirling *et al*. 2021. *BMC Bioinformatics*). In brief, images were analyzed using a method to identify distinct nuclei and count nuclei in association with either DARPP-32 or GFAP (see Supplemental Note 1). Exported data was further processed to identify “Astrocytes” and “MSN” objects which shared a primary object “Nuclei”. These mixed identity nuclei were counted and added to each population of astrocytes and MSN having distinct identity.

### Statistical Modeling of Somatic Expansion

All statistical tests were performed in RStudio with R v4.5.2. Linear modeling applied the lm() function and linear mixed modeling applied the lmer() fuction.

### Blood Somatic Expansion Models

For blood DNA, log-transformed RSE or FER values were used for downstream calculation of residuals after adjusting for the significant effect of inherited uninterrupted CAG repeat length, age at sampling, and their interaction in multiple linear regression as follows:

log(RSE or FER) ∼ Inherited CAG + Sample Collection Age + Inherited CAG*Sample Collection Age

Addition of the significant effect of CAG-CCG LOI genotype to this model yielded a final model for each blood somatic expansion data set as follows:

log(RSE or FER) ∼ Inherited CAG + Sample Collection Age + Genotype + Inherited CAG*Sample Collection Age

### Bulk Brain Tissue Somatic Expansion Models

In canonicals only, all tissues combined (linear mixed model used for comparing instability between tissues):

log(RSE or FER) ∼ Inherited CAG + Disease Duration + Inherited CAG*Disease Duration + Tissue + Random Variable(donor)

In all donors, each tissue analyzed separately (simple linear model used for comparing between genotypes within a given tissue: Caudate, Putamen, Frontal Cortex and Cerebellum):

Log(RSE or FER) ∼ Inherited CAG + Disease Duration + CAG*Disease Duration + Genotype

### Bulk Caudate Small Pool Expansion Models

Linear mixed modeling of single-molecule somatic CAG expansion in bulk caudate was based on the linear regression model previously applied to RSE and FER, but with expanded CAG repeat length of each molecule observation as the dependent variable and a random variable for each donor to account for multiple observations. The resulting model was significant for inherited uninterrupted CAG repeat length, disease duration, and the interaction of these fixed variables as follows:

CAG Molecule Length ∼ Inherited CAG + Disease Duration + Inherited CAG*Disease Duration, with a random variable by donor

Notably, age at death was a nonsignificant predictor of CAG repeat expansion length in all evaluated caudate models. CAG-CCG LOI genotype was nonsignificant when added as a fixed effect, as follows:

CAG Molecule Length ∼ Inherited CAG + Disease Duration + CAG-CCG LOI + Disease Duration*Inherited CAG, with a random variable by donor

### Caudate MSN Small Pool Expansion Models

When linear mixed modeling was applied to single-molecule somatic CAG expansion in canonical caudate MSNs, expansion was significantly associated with inherited uninterrupted CAG repeat length, disease duration, and the interaction of these two variables as follows:

CAG Molecule Length ∼ Inherited CAG + Disease Duration + CAG*Disease Duration, with a random variable by donor

In the combined MSN data set, the addition of CAG-CCG LOI genotype was highly significant when added as a fixed variable, as follows:

CAG Molecule Length ∼ Inherited CAG + Disease Duration + CAG-CCG LOI + Disease Duration*Inherited CAG, with a random variable by donor

Addition of an interaction between disease duration and CAG-CCG LOI genotype further showed significant association with CAG-CCG LOI genotype, disease duration, as well as the interaction of disease duration with both CAG-CCG LOI genotype and inherited uninterrupted CAG:

CAG Molecule Length ∼ Inherited CAG + Disease Duration + CAG-CCG LOI + Disease Duration*Inherited CAG + Disease Duration*CAG-CCG LOI, with a random variable by donor

