## Supplemental Figures for "Loss of interruption in the *HTT* CAG repeat is associated with increased somatic expansion and loss of medium spiny neurons in HD"

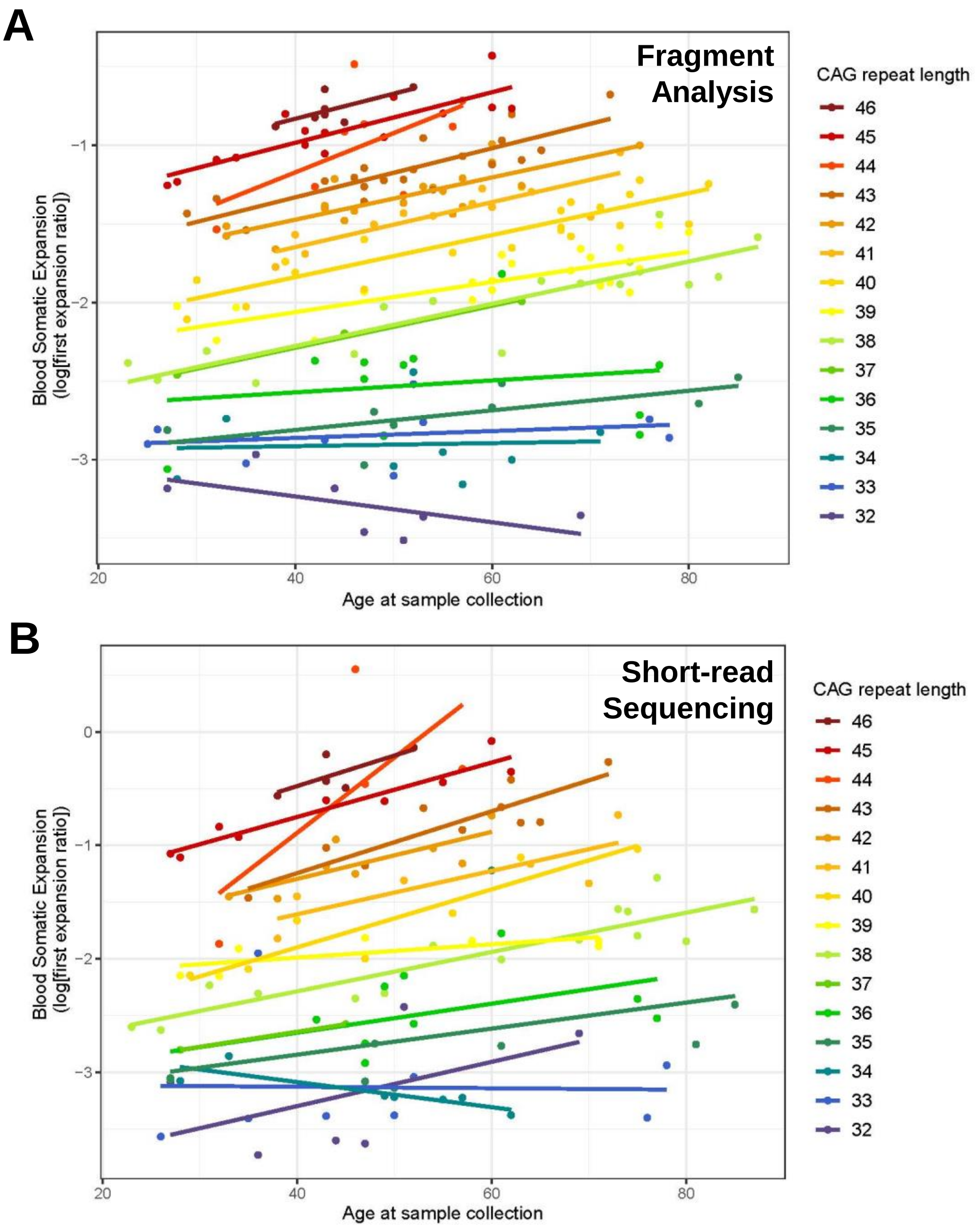


**Supplemental Figure 1. Small somatic expansions increase with inherited uninterrupted CAG repeat length and age at sample collection in blood.** Log-transformed first expansion ratios from (**A)** fragment analysis and (**B**) MiSeq for blood DNA of 191 individuals with canonical alleles and CAG repeat lengths of 32 – 46, split by CAG repeat length and compared based on age at sample collection. CAG repeat lengths > 46 have been excluded due to the small number of individuals at each of these repeat lengths.

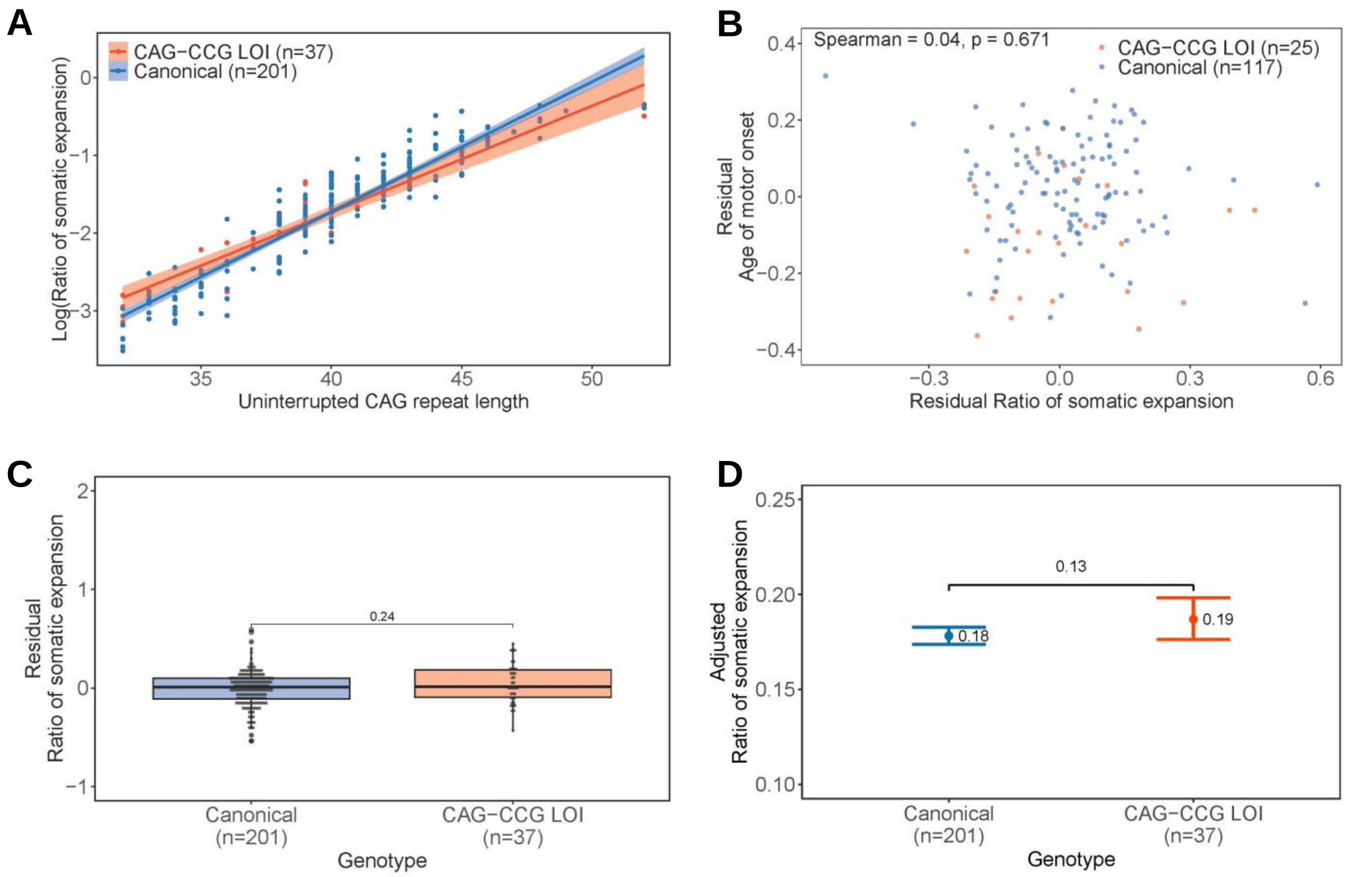


**Supplemental Figure 2. First expansion ratio by fragment analysis increases with inherited uninterrupted CAG repeat length in blood but is unchanged of HD patients with the CAG-CCG LOI variant. (A)** Uncorrected first expansion ratios in canonical and CAG-CCG LOI blood, by uninterrupted CAG repeat length. Log-transformed first expansion ratios from blood DNA of 201 canonical and 37 CAG-CCG LOI variant individuals, compared according to matched uninterrupted CAG repeat lengths. Shaded areas indicate 95% confidence intervals. **(B)** Residual first expansion ratio scores of blood DNA from 117 canonical and 25 CAG-CCG LOI donors with known age at motor onset, compared to residual motor onset scores after accounting for uninterrupted CAG repeat length, showing no association between residual somatic expansion and residual motor onset (Spearman p=0.671). **(C)** Residual first expansion ratio scores of blood DNA from 201 canonical and 37 CAG-CCG LOI donors, after accounting for the effects of uninterrupted CAG repeat length, age at sample collection, and their interaction. Residuals do not differ between genotypes (T-test p=0.24). **(D)** Estimated marginal means plot indicating the effect of genotype on somatic expansion (as measured by first expansion ratio) based on a linear model which accounts for genotype, uninterrupted CAG repeat length, age at blood sample collection, and the interaction between CAG repeat length and sample collection age for all canonical (n=201) and CAG-CCG LOI (n=37) samples with uninterrupted CAG repeat lengths of 32-52, indicating no significant effect of CAG-CCG LOI genotype on first expansion ratios (p=0.13). Error bars indicate 95% confidence intervals.

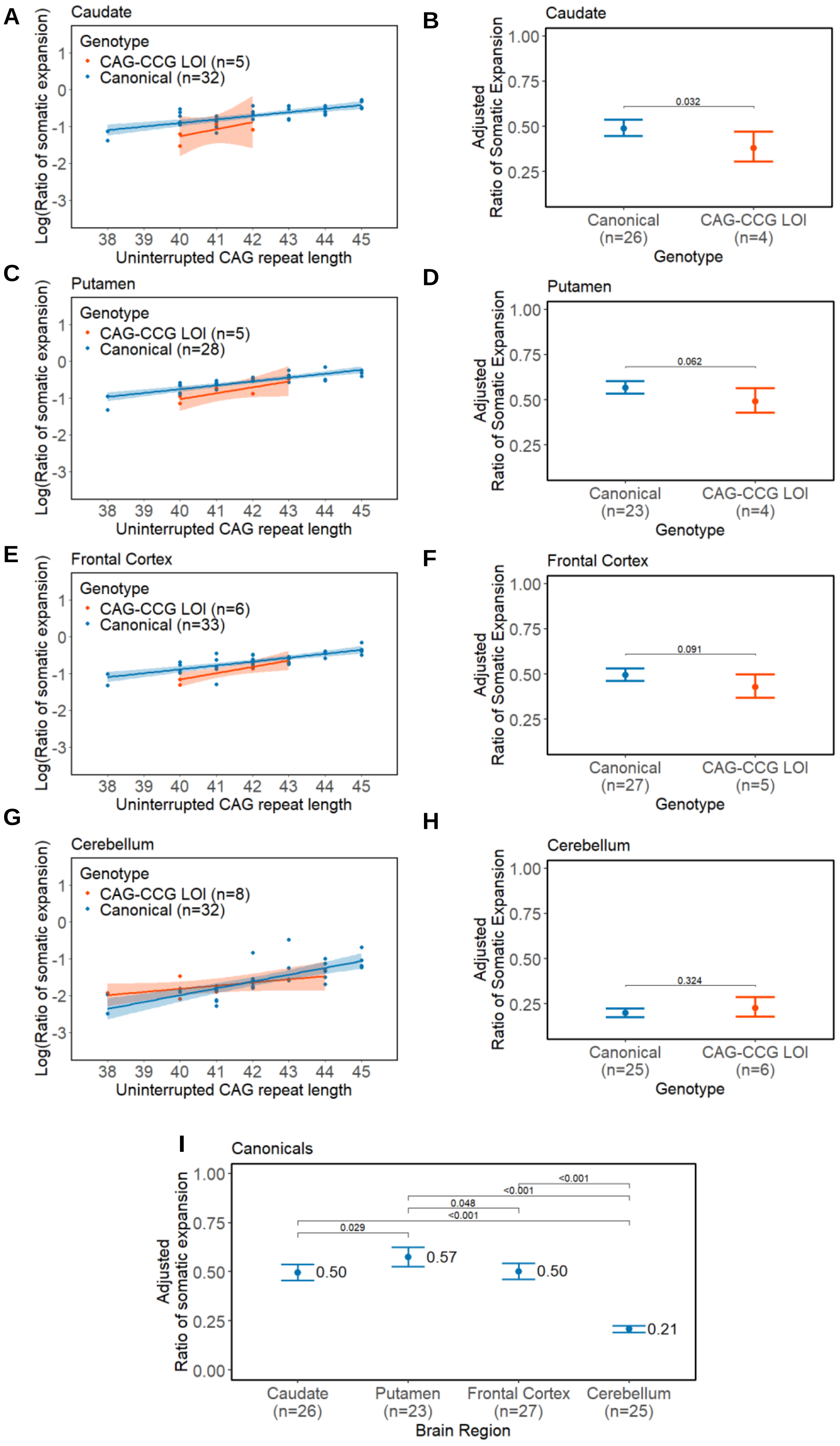

A
Caudate
Genotype
CAG-CCG LOI (n=5)
Canonical (n=32)
Log(Ratio of somatic expansion)
Uninterrupted CAG repeat length
B
Caudate
Adjusted
Ratio of Somatic Expansion
0.032
Canonical
(n=26)
CAG-CCG LOI
(n=4)
Genotype
C
Putamen
Genotype
CAG-CCG LOI (n=5)
Canonical (n=28)
Log(Ratio of somatic expansion)
Uninterrupted CAG repeat length
D
Putamen
Adjusted
Ratio of Somatic Expansion
0.062
Canonical
(n=23)
CAG-CCG LOI
(n=4)
Genotype
E
Frontal Cortex
Genotype
CAG-CCG LOI (n=6)
Canonical (n=33)
Log(Ratio of somatic expansion)
Uninterrupted CAG repeat length
F
Frontal Cortex
Adjusted
Ratio of Somatic Expansion
0.091
Canonical
(n=27)
CAG-CCG LOI
(n=5)
Genotype
G
Cerebellum
Genotype
CAG-CCG LOI (n=8)
Canonical (n=32)
Log(Ratio of somatic expansion)
Uninterrupted CAG repeat length
H
Cerebellum
Adjusted
Ratio of Somatic Expansion
0.324
Canonical
(n=25)
CAG-CCG LOI
(n=6)
Genotype
I
Canonicals
Adjusted
Ratio of somatic expansion
<0.001
<0.001
0.048
<0.001
0.029
0.50
0.57
0.50
0.21
Caudate
(n=26)
Putamen
(n=23)
Frontal Cortex
(n=27)
Cerebellum
(n=25)
Brain Region

**Supplemental Figure 3. The CAG-CCG LOI variant does not increase small somatic expansions in brain tissue of HD patients**. (**A, C, E, G**) The log-transformed fragment analysis first expansion ratio varies by uninterrupted CAG repeat length in brain tissues from canonical and CAG-CCG LOI variant donors (**B, D, F, H**) Estimated marginal means of the first expansion ratio, accounting for genotype, uninterrupted CAG repeat length, disease duration, and the interaction of CAG with disease duration. Significant differences in the adjusted ratio of somatic expansion were observed in the caudate and marginally in the putamen in donors with the CAG-CCG LOI (p-value: 0.032 and 0.062, respectively). Error bars represent 95% confidence intervals. (**I**) The ratio of somatic expansion, adjusted for uninterrupted CAG repeat length and age at donation, differs between caudate, putamen, and frontal cortex relative to the cerebellum in canonical donors ($p<2\times10^{-16}$).

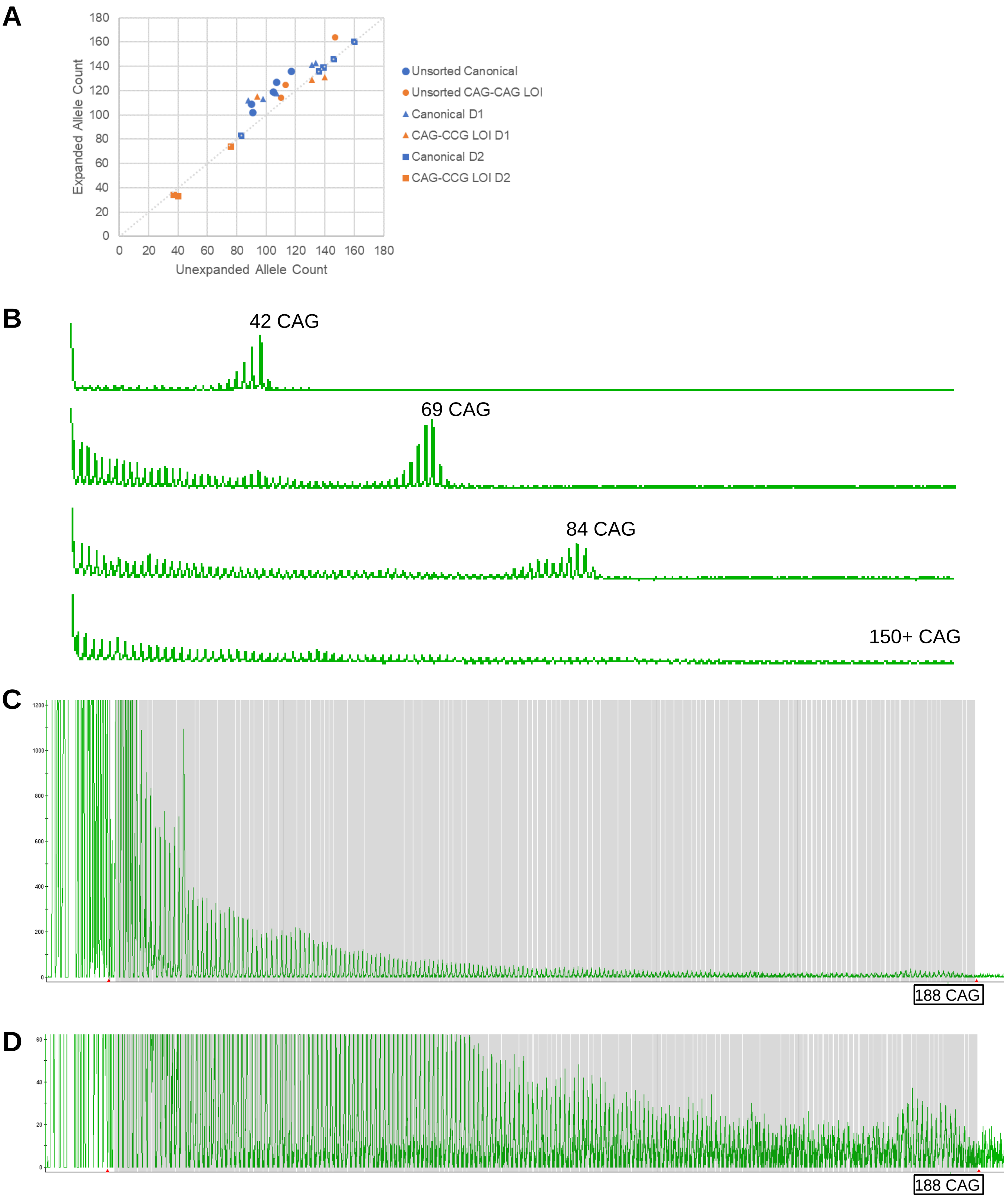


**Supplemental Figure 4. Triplet-primed small pool PCR (TPSP-PCR) analysis of single-molecule somatic CAG expansions. (A)** TPSP-PCR yields approximately equal proportions of CAG molecules from each allele in bulk caudate DNA and sorted caudate MSN DNA. Grey y=x line represents perfectly equal proportions. **(B)** Exemplary TPSP-PCR traces and genotype calls. **(C)** Ascertainment of a very large (>150) CAG repeat expansion at single-molecule resolution by TPSP-PCR at range of 1200rfu and **(D)** 60rfu.

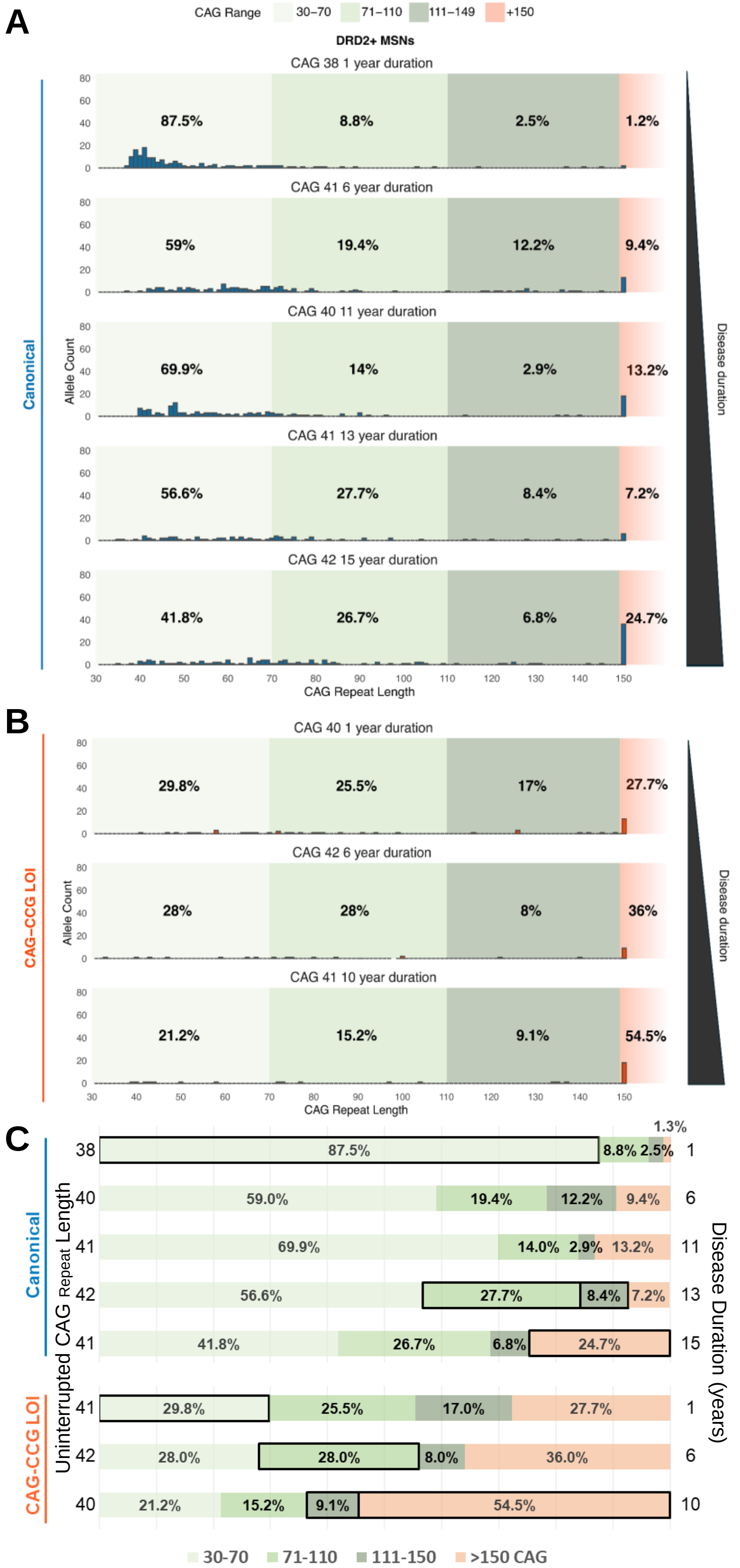


**Supplemental Figure 5. Large somatic expansions in D2 MSNs increase in proportion with disease duration and CAG-CCG LOI genotype.** Single-molecule *HTT* CAG repeat distributions in D2 MSNs from (**A**) canonical-sequence HD donors and (**B**) CAG-CCG LOI variant HD donors. (**C**) Grouped frequencies of small (<50), moderate (51-110), large (111-150), and very large (>150) CAG expansions in D2 MSNs. Proportions boxed in black represent the highest proportion of a given CAG length group among canonical donors or among CAG-CCG LOI donors.
