## Supplemental Tables for "Loss of interruption in the *HTT* CAG repeat is associated with increased somatic expansion and loss of medium spiny neurons in HD"

**Supplemental Table 1.** Multiple linear regression models testing the association between somatic expansion and explanatory variables in peripheral blood and brain tissue using short-read sequencing ratio of somatic expansion (RSE).

| **Blood Somatic Expansion** | | | | | | | |
|---|---|---|---|---|---|---|---|
| No. | Model | Adjusted $r^2$ | *p*-value for model | Parameter values | | | |
| | | | | Sample size | Explanatory variable | Effect in units | *p*-value for explanatory variable |
| 1.* | log(RSE) ~ Inherited CAG + Genotype + Sample Collection Age + Inherited CAG*Sample Collection Age | 0.8924 | <2.2x10$^{-16}$ | 125 Canonical<br>24 CAG-CCG LOI | **Inherited CAG Repeat Length** | **0.224** | **<2x10$^{-16}$** |
| | | | | | **CAG-CCG LOI** | **-0.275** | **6.43x10$^{-4}$** |
| | | | | | **Sample Collection Age** | **0.016** | **2.77x10$^{-13}$** |
| | | | | | **CAG*Sample Collection Age** | **0.001** | **1.93x10$^{-3}$** |
| **Brain Tissue Somatic Expansion (Canonical Donors)** | | | | | | | |
| 2.# | log(RSE) ~ Inherited CAG + Tissue + Disease Duration + Inherited CAG*Disease Duration + Random Variable by Donor | n/a | <2.2x10$^{-16}$ | 27 Canonical | **Inherited CAG Repeat Length** | **0.186** | **0.00281** |
| | | | | | **Caudate** | **1.238** | **<2x10$^{-16}$** |
| | | | | | **Putamen** | **1.579** | **<2x10$^{-16}$** |
| | | | | | **Frontal Cortex** | **1.467** | **<2x10$^{-16}$** |
| | | | | | Disease Duration | 0.002 | 0.86157 |
| | | | | | CAG * Disease Duration | 0.0006 | 0.87328 |
| **Caudate Somatic Expansion (All Donors)** | | | | | | | |
| 3.* | log(RSE) ~ Inherited CAG + Gentoype+ Disease Duration + Inherited CAG*Disease Duration | 0.484 | 1.45x10$^{-3}$ | 20 Canonical<br>5 CAG-CCG LOI | **Inherited CAG Repeat Length** | **0.211** | **0.0454** |
| | | | | | **CAG-CCG LOI** | **-0.482** | 0.0560 |
| | | | | | Disease Duration | 0.0005 | 0.9719 |
| | | | | | CAG* Disease Duration | -0.002 | 0.7503 |
| **Putamen Somatic Expansion (All Donors)** | | | | | | | |
| 4.* | log(RSE) ~ Inherited CAG + Genotype + Disease Duration + Inherited CAG*Disease Duration | 0.7617 | 3.353x10$^{-6}$ | 18 Canonical<br>5 CAG-CCG LOI | **Inherited CAG Repeat Length** | **0.243** | **0.000789** |
| | | | | | **CAG-CCG LOI** | **-0.360** | **0.026230** |
| | | | | | Disease Duration | 0.007 | 0.434376 |
| | | | | | CAG* Disease Duration | -0.003 | 0.509837 |
| **Frontal Cortex Somatic Expansion (All Donors)** | | | | | | | |
| 5.* | log(RSE) ~ Inherited CAG + Genotype + Disease Duration + Inherited CAG*Disease Duration | 0.7336 | 1.361x10$^{-6}$ | 20 Canonical<br>6 CAG-CCG LOI | **Inherited CAG Repeat Length** | **0.177** | **0.0215** |
| | | | | | CAG-CCG LOI | -0.264 | 0.0815 |

* - Canonical alleles were used as the reference for the CAG-CCG LOI explanitory variable.

\# - The Cerebellum was used as the reference for the Tissue explanitory variable.

| | | | | | | | |
|---|---|---|---|---|---|---|---|
| | | | | | Disease Duration | 0.018 | 0.1008 |
| | | | | | CAG* Disease Duration | 0.005 | 0.4413 |
| **Cerebellum Somatic Expansion (All Donors)** | | | | | | | |
| 6.* | log(RSE) ~ Inherited CAG + Genotype + Disease Duration + Inherited CAG*Disease Duration | 0.2557 | 0.0214 | 32 Canonical<br>7 CAG-CCG LOI | Inherited CAG Repeat Length | 0.045 | 0.659 |
| | | | | | CAG-CCG LOI | -0.181 | 0.517 |
| | | | | | Disease Duration | -0.002 | 0.924 |
| | | | | | CAG* Disease Duration | 0.008 | 0.217 |

* - Canonical alleles were used as the reference for the CAG-CCG LOI explanitory variable.
### -  The Cerebellum was used as the reference for the Tissue explanitory variable.

**Supplemental Table 2.** Multiple linear regression models testing the association between somatic expansion and explanatory variables in peripheral blood and brain tissue using fragment analysis first expansion ratio (FER).

| **Blood Somatic Expansion** | | | | | | | |
|---|---|---|---|---|---|---|---|
| No. | Model | Adjusted $r^2$ | *p*-value for model | Parameter values | | | |
| | | | | Sample size | Explanatory variable | Effect in units | *p*-value for explanatory variable |
| 1.* | log(FER) ~ Inherited CAG + Genotype + Sample Collection Age + Inherited CAG*Sample Collection Age | 0.9427 | $<2x10^{-16}$ | 201 Canonical 37 CAG-CCG LOI | **Inherited CAG Repeat Length** | **0.1852** | **$<2x10^{-16}$** |
| | | | | | CAG-CCG LOI | 0.0482 | 0.13 |
| | | | | | **Sample Collection Age** | **0.0121** | **$<2x10^{-16}$** |
| | | | | | **CAG*Sample Collection Age** | **0.0016** | **$7.39\ x10^{-15}$** |
| **Brain Tissue Somatic Expansion (Canonical Donors)** | | | | | | | |
| 2.# | log(FER) ~ Inherited CAG + Tissue + Disease Duration + Inherited CAG*Disease Duration + Random Effect (Donor) | n/a | $<2x10^{-16}$ | 28 donors | **Inherited CAG Repeat Length** | **0.1149** | **0.0001** |
| | | | | | **Caudate** | **0.8670** | **$<2x10^{-16}$** |
| | | | | | **Putamen** | **1.0115** | **$<2x10^{-16}$** |
| | | | | | **Frontal Cortex** | **0.8777** | **$<2x10^{-16}$** |
| | | | | | Disease Duration | 0.0069 | 0.1232 |
| | | | | | CAG * Disease Duration | -0.0004 | 0.8034 |
| **Caudate Somatic Expansion (All Donors)** | | | | | | | |
| 3.* | log(FER) ~ Inherited CAG + Genotype + Disease Duration + Inherited CAG*Disease Duration | 0.5610 | $4.6x10^{-5}$ | 26 Canonical 4 CAG-CCG LOI | **Inherited CAG Repeat Length** | **0.1013** | **0.0140** |
| | | | | | **CAG-CCG LOI** | **-0.2541** | **0.0323** |
| | | | | | Disease Duration | 0.0085 | 0.1897 |
| | | | | | CAG* Disease Duration | -0.0015 | 0.5789 |
| **Putamen Somatic Expansion (All Donors)** | | | | | | | |
| 4.* | log(FER) ~ Inherited CAG + Genotype + Disease Duration + Inherited CAG*Disease Duration | 0.7921 | $5.0x10^{-8}$ | 23 Canonical 4 CAG-CCG LOI | **Inherited CAG Repeat Length** | **0.1181** | **$6.78x10^{-5}$** |
| | | | | | CAG-CCG LOI | -0.1462 | 0.0620 |
| | | | | | **Disease Duration** | **0.0093** | **0.0343** |
| | | | | | CAG* Disease Duration | -0.0020 | 0.2545 |

* - Canonical alleles were used as the reference for the CAG-CCG LOI explanatory variable.

### - The Cerebellum was used as the reference for the Tissue explanatory variable.

| | | | | | | | |
|---|---|---|---|---|---|---|---|
| **Frontal Cortex Somatic Expansion (All Donors)** | | | | | | | |
| 5.* | log(FER) ~ Inherited CAG + Genotype + Disease Duration + Inherited CAG*Disease Duration | 0.6640 | $6.5 \times 10^{-7}$ | 27 Canonical<br>5 CAG-CCG LOI | **Inherited CAG Repeat Length** | **0.1186** | **0.0003** |
| | | | | | CAG-CCG LOI | -0.1425 | 0.0905 |
| | | | | | Disease Duration | 0.0046 | 0.3498 |
| | | | | | CAG* Disease Duration | -0.0015 | 0.4839 |
| **Cerebellum Somatic Expansion (All Donors)** | | | | | | | |
| 6.* | log(FER) ~ Inherited CAG + Genotype + Disease Duration + Inherited CAG*Disease Duration | 0.5801 | $1.82 \times 10^{-5}$ | 25 Canonical<br>6 CAG-CCG LOI | **Inherited CAG Repeat Length** | **0.1355** | **0.0085** |
| | | | | | CAG-CCG LOI | 0.1299 | 0.3238 |
| | | | | | Disease Duration | 0.0055 | 0.5030 |
| | | | | | CAG* Disease Duration | 0.0010 | 0.7475 |

* - Canonical alleles were used as the reference for the CAG-CCG LOI explanatory variable.

### - The Cerebellum was used as the reference for the Tissue explanatory variable.

#### Supplemental Table 3. Caudate Donors

**Caudate Donors for D1/D2 FANS and TPSP-PCR**

| Case No. | CAG-CCG Sequence | Diag CAG | Unint CAG | Motor Onset (years) | Psych Onset (years) | Age at Death (years) | Disease Duration (years) | CAP100 | HD grade | PMD (hours) | Cause of Death |
|---|---|---|---|---|---|---|---|---|---|---|---|
| **HDB-250** | Canonical | 17/38 | 38 | 72 | 72 | 73 | 1 | 90.0 | 1 | 5 | MAID |
| **HBD-062** | Canonical | 15/41 | 41 | 61 | 59 | 67 | 6 | 113.6 | 2 | unk | Unavailable |
| **HBD-240** | Canonical | 15/40 | 40 | 64 | 64 | 75 | 11 | 115.6 | 1 | 24 | MAID |
| **HDB-242** | Canonical | 17/41 | 41 | 55 | 52 | 68 | 13 | 115.3 | 2 | 15 | Complications of HD, bronchopneumonia and pulmonary embolism |
| **HDB-159** | Canonical | 17/42 | 42 | 54 | 59 | 69 | 15 | 127.6 | 2 | 7 | Unavailable |
| **HC086** | CAG-CCG LOI | 17/38 | 40 | 45 | Unk | 46 | 1 | 70.9 | 1 | 18 | Head injury |
| **HC103** | CAG-CCG LOI | 19/40 | 42 | 35 | 40 | 41 | 6 | 75.8 | 1 | 11 | Renal failure |
| **HC149** | CAG-CCG LOI | 20/39 | 41 | 44 | 42 | 54 | 10 | 91.5 | 2 | 6.5 | Pneumonia |

**Caudate Donors for DARPP-32+ MSN Counting**

| Case No. | CAG-CCG Sequence | Diag CAG | Unint CAG | Motor Onset (years) | Psych Onset (years) | Age at Death (years) | Disease Duration (years) | CAP100 | HD grade | PMD (hours) | Cause of Death |
|---|---|---|---|---|---|---|---|---|---|---|---|
| **COB-064** | Canonical | 17/19 | 19 | n/a | n/a | 86 | n/a | n/a | n/a | 17 | Unavailable |
| **HDB-258** | Canonical | control | control | n/a | n/a | 72 | n/a | n/a | n/a | 24 | Metastatic breast cancer |
| **HDB-260** | Canonical | control | control | n/a | n/a | 62 | n/a | n/a | n/a | 12 | Graft-versus-host disease of intestines |
| **HDB-211** | Canonical | 22/44 | 44 | 49 | Unk | 58 | 9 | 125.1 | 1 | 8 | Unavailable |
| **HDB-240** | Canonical | 15/40 | 40 | 64 | 64 | 75 | 11 | 115.6 | 1 | 24 | MAID |
| **HDB-250** | Canonical | 17/38 | 38 | 72 | 72 | 73 | 1 | 90.0 | 1 | 5 | MAID |
| **HDB-181** | Canonical | 17/43 | 43 | 45 | 38 | 62 | 17 | 124.2 | ~2 | 13 | Aspiration pneumonia |
| **HDB-182** | Canonical | 17/41 | 41 | 62 | 56 | 76 | 14 | 128.8 | ~2 | 24 | Unavailable |
| **HDB-212** | Canonical | 17/42 | 42 | 45 | Unk | 66 | 21 | 122.0 | 3 | 22 | HD, unspecified complications |
| **HDB-242** | Canonical | 17/41 | 41 | 55 | 52 | 68 | 13 | 115.3 | 2 | 15 | Complications of HD, bronchopneumonia and pulmonary embolism |
| **HDB-259** | Canonical | 15/42 | 42 | 51 | 58 | 65 | 14 | 120.2 | 2 | 24 | Unavailable |
| **HDB-179** | Canonical | 23/43 | 43 | 54 | Unk | 78 | 24 | 156.2 | 3 | 7 | HD, unspecified complications |
| **HDB-180** | Canonical | 17/42 | 42 | 46 | 46 | 75 | 29 | 138.7 | 3 | Unk | HD, unspecified complications |
| **HDB-185** | Canonical | 17/45 | 45 | 41 | 42 | 64 | 23 | 147.9 | 3 | 20 | Bronchopneumonia |
| **HDB-178** | Canonical | 19/44 | 44 | 48 | 43 | 68 | 20 | 146.7 | 4 | 14 | Aspiration pneumonia |
| **HC103** | CAG-CCG LOI | 19/40 | 42 | 35 | 40 | 41 | 6 | 75.8 | 1 | 11 | Renal failure |
| **HC095** | CAG-CCG LOI | 20/39 | 41 | 40 | Unk | 66 | 26 | 111.9 | 2 | 12 | HD, unspecified complications |
| **HC149** | CAG-CCG LOI | 20/39 | 41 | 44 | 42 | 54 | 10 | 91.5 | 2 | 6.5 | Pneumonia |
| **HC176** | CAG-CCG LOI | 21/36 | 38 | 65 | Unk | 86 | 21 | 106.0 | 3 | 7.5 | Unavailable |

**Supplemental Table 4.** Multiple linear regression models testing the association between somatic expansion and explanatory variables in bulk caudate tissue and and FANS-sorted medium spiny neurons by TPSP-PCR.

| **Bulk Caudate Somatic Expansion (All donors)** | | | | | | |
|---|---|---|---|---|---|---|
| No. | Model | *p*-value for model | Parameter values | | | |
| | | | Sample size (CAG molecules from expanded HD allele) | Explanatory variable | Effect in units | *p*-value for explanatory variable |
| 1.* | log(RSE) ~ Inherited CAG + Genotype + Sample Collection Age + Inherited CAG*Sample Collection Age | 0.00384 | 996 (8 donors) | **Inherited CAG Repeat Length** | **2.637** | **0.00952** |
| | | | | CAG-CCG LOI | -0.757 | 0.7898 |
| | | | | Disease Duration | 0.005 | 0.9817 |
| | | | | CAG*Disease Duration | -0.231 | 0.0430 |
| | | | | CAG-CCG LOI*Disease Duration | 0.234 | 0.4257 |
| **Medium Spiny Neuron Somatic Expansion (Canonical Donors)** | | | | | | |
| 2. | CAG Molecule Length ~ Inherited CAG + Disease Duration + CAG*Disease Duration + Random Variable by Donor | $1.069 \times 10^{-4}$ | 627 (5 donors) | **Inherited CAG Repeat Length** | 4.702 | **0.0235** |
| | | | | Disease Duration | 0.506 | 0.2694 |
| | | | | **CAG*Disease Duration** | 0.428 | **0.0322** |
| **Medium Spiny Neuron Somatic Expansion (All Donors)** | | | | | | |
| 3.* | CAG Molecule Length ~ Inherited CAG + Disease Duration + Genotype + Inherited CAG*Disease Duration + Genotype* Disease Duration + Random Variable by Donor | $2.871 \times 10^{-10}$ | 1002 (8 donors) | Inherited CAG Repeat Length | 2.856 | 0.1487 |
| | | | | Disease Duration | 0.880 | 0.0509 |
| | | | | **CAG-CCG LOI** | 35.156 | **$2.18 \times 10^{-10}$** |
| | | | | **CAG*Disease Duration** | 0.458 | **0.0393** |
| | | | | **CAG-CCG LOI*Disease Duration** | 1.420 | **0.0156** |

* - Canonical alleles were used as the reference for the CAG-CCG LOI explanitory variable.
